# A Light Sensitive Hyaluronic Acid-based Hydrogel for 3D Culturing, Differentiation, and Harvesting of Mesenchymal Stromal Cells

**DOI:** 10.1101/2024.08.13.607705

**Authors:** Francesco Palmieri, Marieme Gueye, Lucia Vicario del Rio, Saskia Bunschuh, Pradeep Chopra, Silvia Mihăilă, Tina Vermonden, Riccardo Levato, Geert-Jan Boons

**Author notes:** **Corresponding Author Geert-Jan Boons** – Division of Chemical Biology and Drug Discovery, Utrecht Institute for Pharmaceutical Sciences, Utrecht University, Universiteitsweg 99, Utrecht 3584 CG, The Netherlands; Complex Carbohydrate Research Center, University of Georgia, 315 Riverbend Road, Athens, Georgia 30602, United States; and.

## Abstract

Hydrogels derived from natural sources are commonly used for 3D cell culturing due to their favourable interactions with cells. However, these biomaterials (*<u>e</u>*.*<u>g</u>*. Matrigel) suffer from variations in composition, limited mechanical tunability, the presence of xenogenic components and difficulties of cell retrieval. Semi-synthetic hydrogels are emerging to address these limitations, making these attractive for drug delivery and tissue engineering. Here, we describe a hydrogel platform based on hyaluronic acid (HA) modified by (1R,8S,9S)-bicycle[6.1.0]non-4-yn-9-ylmethanol (BCN) and a cross-linker composed of light sensitive *o*-nitrobenzyl moieties and polyethylene glycol (PEG) chains terminating in azides. The two components can undergo strain-promoted azide-alkyne cycloaddition (SPAAC) resulting in rapid gel formation. The stiffness of the hydrogel can be modulated by varying the crosslinker ratio and multi-functionalizing is possible by the incorporation of chemical cues modified by an azide. The fast hydrogelation enables easy encapsulation of cells and harvesting with preservation of viability is possible be short exposure to UV light. Moreover, it is demonstrated that visible light (405 nm) can soften the hydrogel resulting in phenotypical changes of human Mesenchymal Stromal Cells (hMSCs) co-cultured with endothelial colony-forming cells (ECFCs) with high viability. Our findings highlight that the photosensitive HA-based hydrogel provides a versatile and biocompatible platform for cell culture and tissue engineering applications, offering advantages over traditional 3D cell culturing platforms.

## INTRODUCTION

Hydrogels derived from various natural sources are commonly used for 3D cell culturing, offering a supportive environment for cell growth and interaction.^1-5^ However, these hydrogels often suffer from shortcomings and for example may exhibit variable mechanical and biochemical properties due to their complex composition and batch-to-batch variations.^1-3^ This variability can lead to challenges in reproducibility and interpretation of experimental results. Additionally, the tunability of hydrogel properties such as spatial arrangement of molecular signals and mechanical properties may be limited, hindering precise control over cell fate and behavior.^1, 4^ Furthermore, some hydrogels may contain xenogenic components, rendering these unsuitable for therapeutic cell manufacturing applications.^5^

Semi-synthetic hydrogels that can be manufactured in a reproducible manner and that can be designed to precisely tune their mechanical and chemical properties, are emerging as attractive alternatives. Ideally, such a hydrogels should have high water absorption capacity, biocompatibility, and tunable physicochemical properties.^6^ These properties make hydrogels suitable for use as biosensors, drug delivery systems, and matrices for tissue engineering.^7^

Hydrogel fabrication from naturally derived materials is advantageous for biomedical applications due to their innate cellular interactions and cellular-mediated biodegradation.^8-9^ Moreover, a hydrogel should allow for modifications with biological cues, such as cell adhesion peptides to provide further possibilities to control cell fate.

Stimulus-responsive hydrogels that can undergo phase transitions make it possible to embed and harvest cells without the need to use aggressive enzymes to degrade the extracellular matrix that may result in cellular membrane damage, such as depletion or degradation of integrins, cadherins and surface markers, potentially affecting cell signaling and adhesion, differentiation and functional properties. Such hydrogels provide opportunities for regenerative medicine and facilitate harvesting and subsequent examination of phenotypic properties of cultured cells. Temperature responsive hydrogels have been described that exhibit either an increase or decrease in viscosity when raising the temperature.^10-11^ These transitions are generally reversable, and as a result these hydrogels can undergo multiple cycles of sol-gel transitions.^12^ Nevertheless, majority of mammalian cells are able to function normally at 37 °C, producing heat shock proteins (HSPs) at 41 °C ^13^ and changing their lipidic composition to cold stress (*e*.*g*. 32 °C).^14^ Therefore, even small thermal variation can dramatically influence the outcome of biological readout without a major modification in the materials properties. Moreover, the incorporation of chemical entities to influence cell faith can influence thermal responsiveness making it more difficult to develop multifunctional responsive materials.

Sol-gel transitions have also been accomplished by pH-responsive hydrogels.^15^ One approach is based on reversible covalent bonds between primary amines and aldehydes or ketones. The pH range required for the sol-gel transitions is, however, not compatible with cell viability. Recent advances have made it possible to induce sol-gel transitions at physiological pH (6.5-7.4) by protonation of specific functional groups, enabling ionic interactions during gelation.^16^ Although this represents an attractive strategy, even slight variations in pH can induce substantial changes in cell viability.^17^

A particularly attractive approach to induce a gel-sol transition is based on hydrogels that contain a photosensitive moiety. Light can then be employed to cleave the hydrogel bundles at predesigned locations in a spatially and temporally controlled manner. Although several photosensitive chemical moieties have been explored such as coumarin derivatives^18^ and ruthenium complexes,^19^ *o*-nitrobenzyl derivatives (*o*-NB) are one the most attractive photolabile groups for biomedical applications,^20-22^ because of straightforward synthesis, easy handling and relative high quantum yields in the photoreaction.^23^ Their one photon activation is possible using a near UV light at 365 nm but absorption profile can been adjusted by modifying the aromatic structure.^24^ UV light, including 365 nm light, can be harmful to cells due to its ability to induce DNA damage, oxidative stress, and other cellular changes.^25-27^ Prolonged or intense exposure to UV light can lead to cell death or other adverse effects. However, it’s important to note that 365 nm light falls within the UVA range of UV light, which is considered less damaging than shorter wavelengths such as UVB and UVC. Therefore short exposure time and low light intensity is essential to avoid cytotoxicity.^27^

We describe here a hyaluronic acid (HA) based hydrogel that includes an *o*-nitrobenzyl carbamate containing cross-linker that can undergo very fast gel formation and degradation and is attractive for harvesting cultured cells. Moreover, we demonstrate that by varying in the light frequency, mechanical properties of the hydrogel can be modulated leading to phenotypical changes of cultured cells. HA is a glycosaminoglycan found in the extracellular matrix of many tissue types (Fig. 1). It is an attractive biopolymer for tissue engineering and regenerative medicine because it is cytocompatibility and can be produced on large scale.^28-30^ It is composed of repeating disaccharides of glucuronic acid (GlcA) and *N*-acetyl glucosamine (GlcNAc). The carboxylic acid of GlcA provides attractive chemical handle to attach various chemical entities by ester or amide formation. We choose to modify HA with (1R,8S,9S)-bicycle[6.1.0]non-4-yn-9-ylmethanol (BCN) to yield polymer **1** that can undergo strain-promoted azide-alkyne cycloaddition (SPAAC) with azides. BCN was selected as strained cyclo-octyne because it exhibits relative low hydrophobicity yet maintains high reactivity with organic azides.^31^ As cross-linker, we employed compound **2**, which incorporates two *o*-nitrobenzyl moieties that are linked through a carbamate linkage to a polyethylene glycol (PEG) chains terminating in azides.^31^ The carbamate linker has greater stability compared to previously employed ester linked derivatives^32^ and also increases the photo-reactivity.^32^ Addition of crosslinker **2** to a solution of functionalized HA **1** resulted in fast hydrogel formation and by employing different molar ratios of crosslinker **2**, the stiffness could be modulated. Short exposure (1 min) to low intensity UV light resulted in complete degradation of the hydrogel. To demonstrate that multifunctional hydrogels can be prepared, integrin derived peptide **3** modified by azide was synthesized and a functionalized hydrogel could be formed by the addition of **2** and **3** to a solution of **1**. hMSCs and endothelial colony forming cell (ECFCs) were embedded in the hydrogel and cell stretching was observed when the hydrogel gel was exposed to visible light to soften it. As a control, a hydrogel was formed using **4** as cross linker that has Matrix Metalloprotease (MMP2) cleavable amino acid sequence.^33^ hMSCs embedded in the hydrogel also exhibited a stretched morphology, which likely is due to partial degradation and softening by secreted MMP2. Almost quantitative retrieval of the encapsulated hMSCs was possible by short UV-mediated photodegradation, which had almost no impact on viability, morphology and adhesive capabilities.

**Figure 1.**
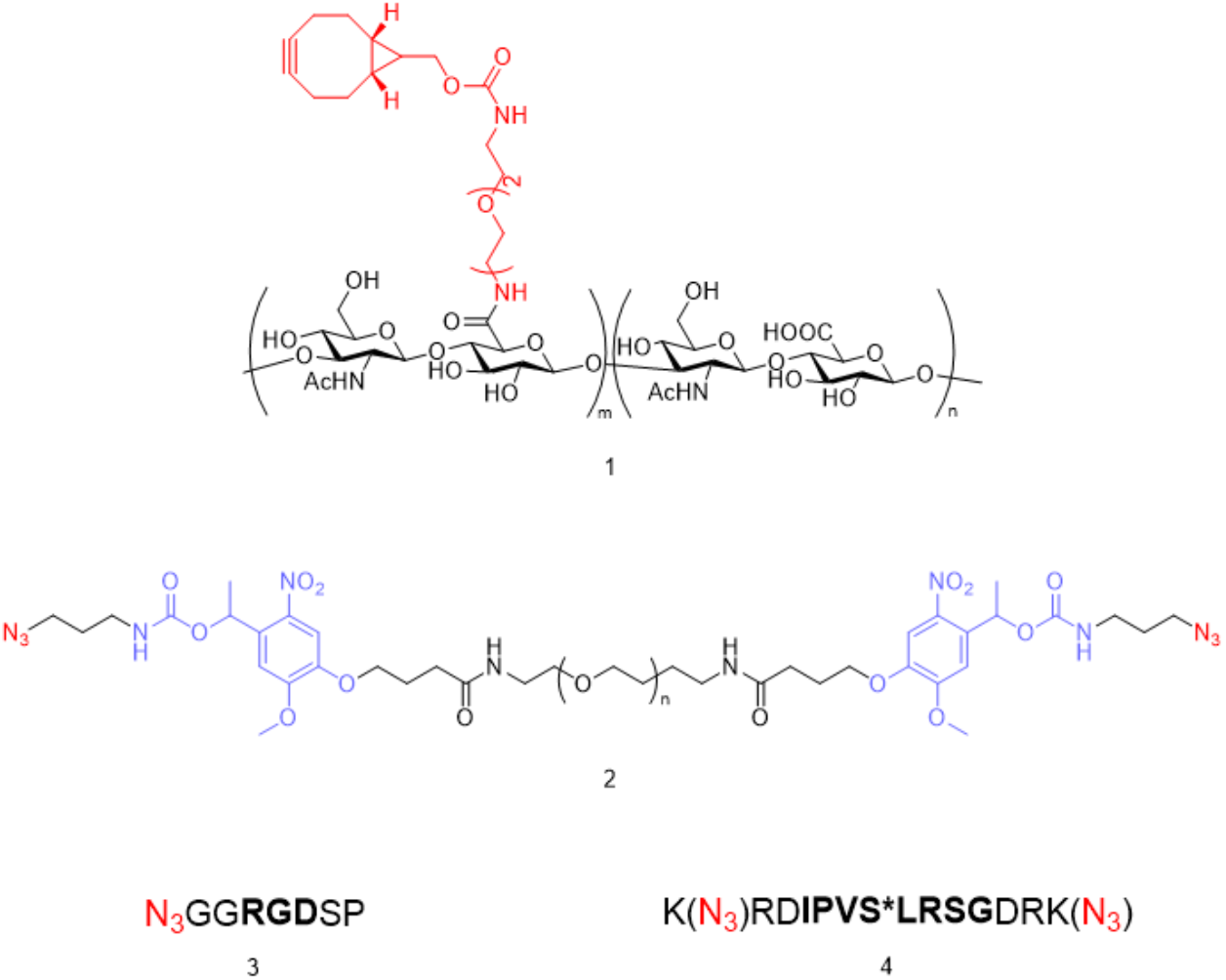
Component for formation of a multifunctional photosensitive hydrogels.

## EXPERIMENTAL SECTION

### SPPS of Adhesive Peptide and MMP2 Cleavable Crosslinker

RGD peptide (compound **3**) and MMP2 cleavable crosslinker (compound **4**) were synthetized by standard Solid Phase Peptide Synthesis (SPPS) using liberty blue peptide synthesizer. In case of peptide **3** Fmoc-Ser-OH was coupled to Rink amide resin, then coupled to the subsequent regular amino acid to build the sequence. The azide moiety was introduced by couple the last amino acid of the sequence to azidoacetic acid. In case of peptide **4**, Fmoc-Lys(N_3_)-OH was coupled to Rink amide resin, then the subsequent regular amino acids were coupled using standard protocols. The N-terminal part of the peptide includes a second Fmoc-Lys(N_3_)-OH moiety which capped by standard capping procedure using acetic anhydride.

### Hydrogel Preparation

HA-BCN was dissolved in PBS (phosphate buffered saline) pH 7.4, to obtain a solution with a final concentration of 5% w/v. The photosensitive crosslinker (compound **2**) was dissolved in milliQ water with a final concentration of 100 mM. Combining these two solutions in a ratio (v/v) of 100/9, 100/6, and 100/3 resulted in the formation of stable hydrogels having a crosslinking degree of 100%, 66%, and 33%, respectively referring to the theoretical BCN units consumed during the crosslinking.

### Mechanical Characterization

The mechanical properties of the hydrogels were analyzed with a rheometer (Discovery HR2, TA Instruments, New Castle, DE, USA) using a cone-plate geometry (20 mm – 1°) performing a time/sweep at 37 °C with strain of 2% and an angular frequency of 10%. The frequency sweep was performed using a cone-plate geometry (20 mm – 1°) performing at 37 °C with strain of 2% and a frequency from 0.1 to 100 Hz. The amplitude sweep was performed using a cone-plate geometry (20 mm – 1°) at 37 °C with a frequency of 2 Hz and a strain from 0.1% to 100%. The photodegradation was analysed with a rheometer (Discovery HR2, TA Instruments, New Castle, DE, USA) using a plate-plate geometry (20 mm – 1°) performing a time/sweep at 37 °C with strain of 2% and an angular frequency of 10% using a UV Lamp (BluePoint UVC extended range) with a filter 320-500 nm, having a light intensity of 230 mW/cm2.

### Cell Culture

Human adipose-derived mesenchymal stromal cells (MSCs), kindly provided by Radboud Institute of Molecular Materials, were cultured using MSC medium at 37 °C in 5% CO_2_. MSC medium was composed of 5 g Alpha MEM (Minimum Essential Medium Alpha Medium, ThermoFisher Scientific, Gibco, #12000063), 1.1 g NaHCO3 (sodium bicarbonate, Sigma Aldrich, #S5761) dissolved into 500 mL demi water. The medium was sterilized by using Stericup Quick Release-GP sterile vacuum filtration system (#S2GPU05RE), supplemented with 10% fetal calf serum (Biowest, #S1810-500, Batch: S00FI) and 1% Penicillin Streptomycin (Pen Strep, Gibco Fisher Scientific, 10000U/ml, #15140122). Cells were passaged up to passage number 10 using 0.05% Trypsin-EDTA (Gibco, #25300-062) for cell detachment and medium was changed every three days. MSCs were counted manually using Trypan Blue solution (Sigma, #T8154). To ensure single cell embedding, MSCs were filtered by Falcon® 70 µm Cell Strainer (BD Falcon, #734-0003) post harvesting.

Endothelial colony-forming cells (ECFCs) were isolated from umbilical cord blood after informed consent was obtained from the mothers. Cells were genetically modified to express Green Fluorescent Protein (GFP). GFP-ECFCs were expanded in EGM-2 medium (Lonza), composed of Endothelial Cell Basal Medium-2 (EBM-2) and supplemented with 10% FBS and growth factors (1% Pen/Strep, 0.4% hFGF-β, 0.1% ascorbic acid, 0.1% VEGF, 0.1% hEGF, 0.1% R3-IGF-1, 0.1% heparin and 0.04% hydrocortisone). They were expanded at 37 ºC in 5% v/v CO_2_ humidified atmosphere. The culture medium was changed every 2 days. ECFCs were used for experimentation at passage numbers 14-18 and were passaged using 0.25% Trypsin-EDTA.

### Cell Embedding

Post cell harvesting, 1.5×10^6^ MSCs per mL were resuspended in HA-BCN 5% w/v solution in PBS buffer and subsequently transferred in a disk-shaped mold of 50 µL. After adding 3 µL of crosslinker solution (100 mM in mQ water), the gel was crosslinked at 37 °C for 10 min. The gels with cells were transferred from the mold to a 12- or 48-well plate (for suspension cell culture) and complete media was added to cover the entire gel. Embedded MSCs were further incubated for 24 h at 37 °C with 5% CO_2_.

### UV Cleavage and Re-seeding

Prior UV exposure, hydrogels were transferred into a glass bottom petri dish (FluoroDish, #FD3510-100). A UV lamp (BluePoint Lamp UVC extended range, filter 320-500 nm) was used to expose MSCs in 3D at an intensity of ∼20 mW/cm^2^ in 30 sec UV light intervals. Between the UV-intervals cells were gently washed out of the hydrogel by applying 1-3 mL of prewarmed complete media. This media (including retrieved cells from hydrogel) was collected in canonical tubes until the entire gel was sufficiently cleaved and subsequently centrifuged to remove remains of hydrogels which accumulated on top of the cell pallet post centrifugation (300 rpm for 3 min at room temperature). Cells were plated in 24-well plates (Corning, #3612) and cultured at standard culture conditions for 24, 48, or 72 h.

### Live-Dead Staining

Live-Dead staining was performed within 24 h after cell embedding, with 4 µM Ethidium-Homodimer 1 (EthD-1) (Thermofisher, #E1169) and 2 µM Calcein-AM, cell permeant dye (Thermofisher, #C1430). Gels or 2D cultured cells were washed with PBS and an appropriate amount live-dead solution, diluted in PBS, was added to cover either the entire gel in case of 3D culture or the entire well bottom in case of 2D culture. After 30 min of incubation at 37 °C with 5% CO_2_, gels were washed twice with PBS (for 5 min) and then directly imaged with Nikon eclipse TS2 microscope. Three to six randomly selected areas per sample were taken and live and dead cells were counted manually. The average ratio of vital over total cells per sample was calculated.

### Metabolic Activity

Metabolic activity was assessed with either PrestoBlue cell viability reagent (Thermo Scientific, #A13262) or Alamar Blue assay which are both resazurin-based solution. When applying it to cells, the reagent is modified by the reducing environment of the viable cells and turns fluorescent. The PrestoBlue reagent was diluted 1:10 with complete media and applied to seeded embedded MSCs in hydrogel. After 1 h of incubation at 37 °C with 5% CO_2_, 1 mL of supernatant was transferred in a 96-well plate and fluorescence was read at 560 nm wavelength with a microplate reader GLOMAX discove. Cell cultures were continued after 3x washing with PBS. The PrestoBlue intensity was normalized against blank (complete media) and the amount of DNA, calculated with DNA quantification assay.

### DNA Quantification

PicoGreen dsDNA quantitation kit (Quant-iT™ PicoGreen™ dsDNA Assay Kits, Thermofisher, #P7589) was used for DNA quantification. After cells were washed once with PBS, cell lysis was initiated by applying ultra-pure water. Cells were then scratched from plate surface, vortexed and stored at -20 °C until performing the assay. The protocol of PicoGreen dsDNA quantitation kit was followed. Briefly, a mixture of 29 µL of sample or standard, 71 µL of PicoGreen solution and 100 µL 1X TE were applied in each well of a 96-well white opaque plate. A standard curve was created by using a sample with known number of MSCs. Post 10 min incubation in the dark, fluorescence was measured with excitation of 484/20 nm and emission of 528/20 nm.

### Statistical Analysis

All experiments were performed at least in triplicates if not otherwise stated. Data visualization and statistical analysis were performed using GraphPad Prism version 9.0.0 (GraphPad Software, San Diego, CA, USA). Statistical significance is considered at p<0.05 and was determined by applying appropriate statistical tests. Data are represented as mean ± standard error of the mean (SEM) or as mean ± standard deviation (SD).

## RESULTS AND DISCUSSION

### Synthesis of HA-BCN 1 and Photosensitive Crosslinker 2 and Mechanical Properties of the Resulting Hydrogels

HA (60 kDa) was coupled with compound **5** (0.15 eq relative to carboxylic acids of HA) in phosphate buffer at pH 5.0 using (4-(4,6-dimethoxy-1,3,5-triazin-2-yl)-4-methyl-morpholinium chloride) (DMTMM) as a coupling reagent resulting in approx. 15% functionalization of the carboxylic acids as determined by NMR integration of acetamido signals from HA monomeric unit and the three different signals belonging to BCN (Scheme 1 and NMR S15). The functionalization of HA was kept at ∼15% of the available carboxylic acids as a compromise between solubility and mechanical properties of the hydrogel. At this level of functionalization, sufficient BCN moieties are present to tune the stiffness of the material. HA-BCN was purified by subsequent dialysis in 10% acetonitrile/H_2_O, 100 mM NaHCO_3_ and finally mQ water.

Crosslinker **2** has two azide moieties at the termini of the linear PEG chains that can react with the BCN moieties of HA derivative **1** to form a hydrogel. In addition, it contains photocleavable *o*-NB moieties that are attached to the PEG chains by a carbamate bond. Recently, it was shown that a carbamate modified o-NB exhibits fast photodegradation and has low hydrolysis rates even at high pH.^32^ The *o*-NB moiety rearranges upon exposure of light resulting in the cleavage of the adjacent benzylic bond. The preparation of **2** started from **6** that was obtained by a reported synthetic procedure^32^ (Scheme 1). Compound **6** was coupled with an excess of 3-azidopropan-1-amine **7** to displace the nitrophenyl esters resulting in the formation of compound **8**, which could be isolated in high yield and purity simply by extraction. Compound **8** was dissolved in THF and treated with aqueous LiOH to hydrolyze the ester to give carboxylic acid **9**. The latter compound was coupled with PEG di-amine (**10**, Mw = 2 kDa) in the presence of 1-[bis(dimethylamino)methylene]-1H-1,2,3-triazolo[4,5-b]pyridinium 3-oxide hexafluoro phosphate (HATU) and 1-hydroxy-7-azabenzotriazole (HOAt) in DMF to yield photosensitive crosslinker **2**. Final compound **2** was obtained in a yield of 39% after purification by size exclusion chromatography over Sephadex LH-20 using DCM/MeOH (1:1, v:v) as the eluent system.

**Scheme 1.**
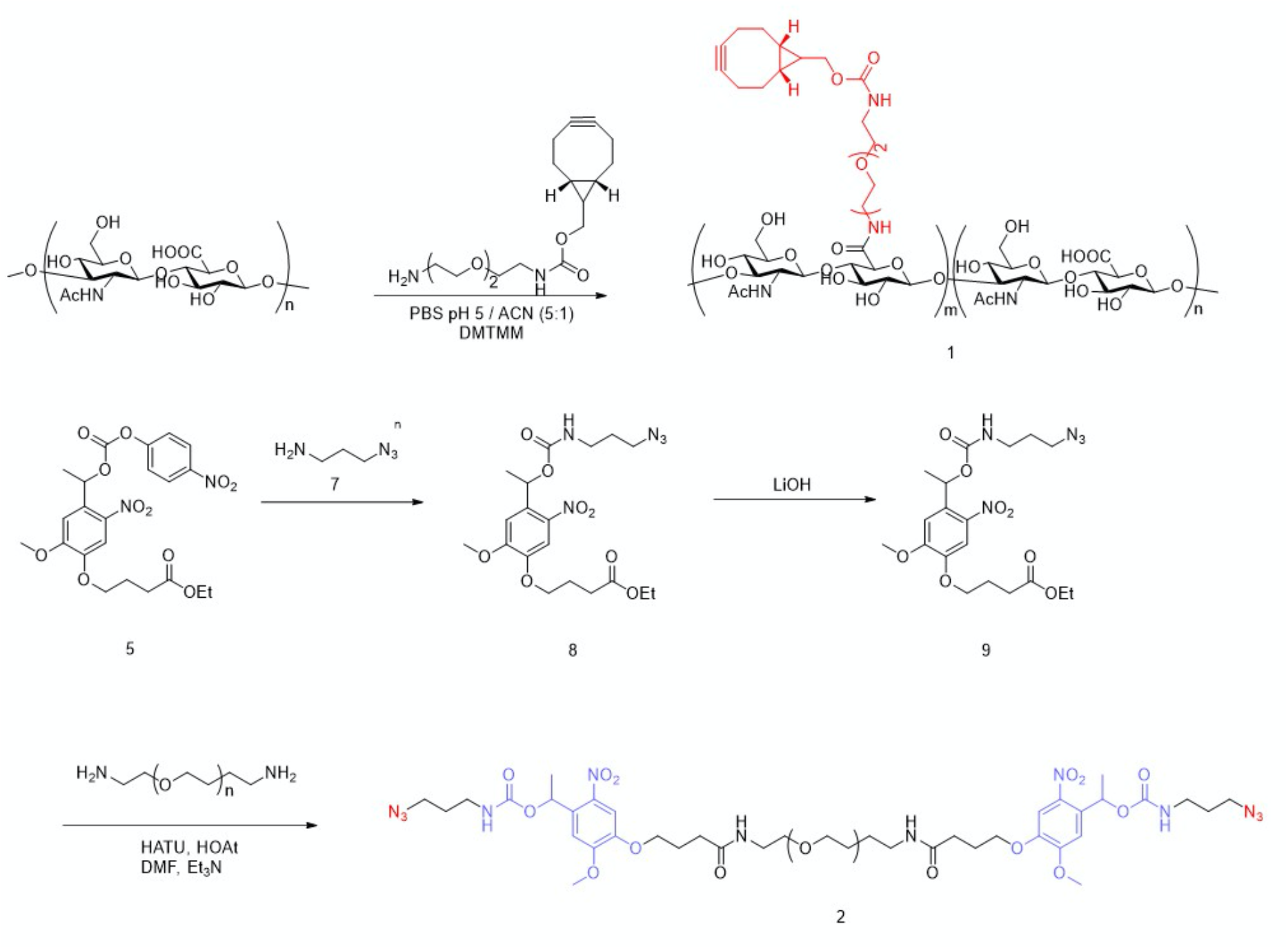
Synthesis of components of modified HA and photosensitive crosslinker.

### Formation and Mechanical Properties of HA Photosensitive Hydrogels

Stiffness of a hydrogel can impact cell fate by influencing properties such adhesion, proliferation differentiation and migration.^34-36^ Hydrogels were formed using different molar ratios of HA-BCN (**1**) and azides in crosslinker (**2**). Rheological properties of these hydrogels were determined by monitoring the storage (G^’^) and the loss (G^’’^) modulus during the formation of the hydrogels (Fig. 2a). Molar ratios of BCN to N_3_ of 1:1, 3:2 and 3:1 gave of G’ of 7.2 kPa, 5.3 kPa and 3.5 kPa, respectively. The increase in stiffness with increasing crosslinker is sinusoidal (Fig. 2b).

**Figure 2.**
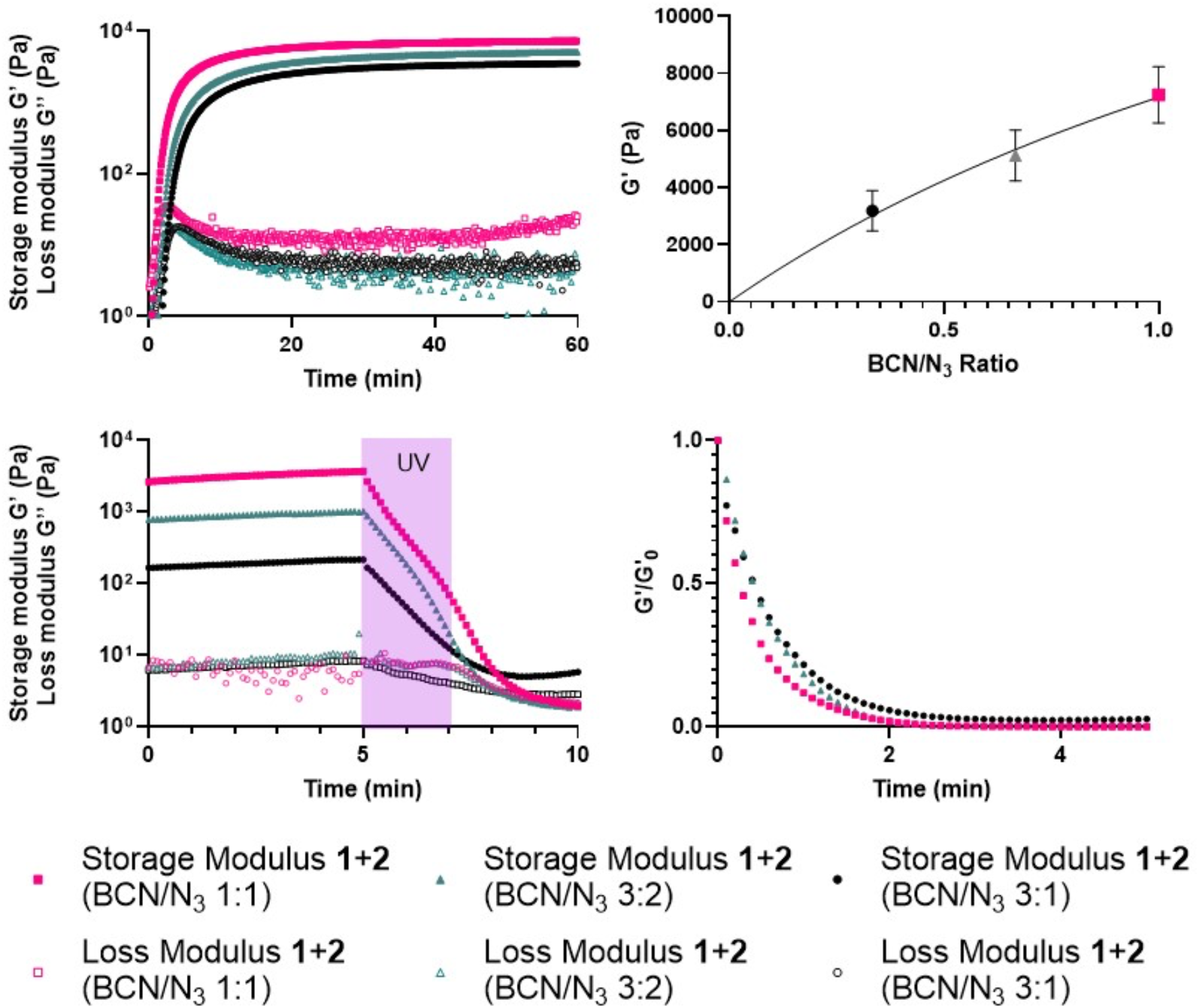
Rheological characterization of photosensitive hydrogels. **a)** Storage (G’) and loss modulus (G’’) at different time points of hydrogel formation by mixing 5% w/v solution of HA-BCN (**1**) with a solution (0.1 M) photocleavable crosslinker (**2**) in a BCN/N_3_ ratio of 1:1 (red squares), 3:2 (gray triangles) and 3:1 (black circle), respectively. **b)** Maximum stiffness of the hydrogels using a ration of BCN/N_3_ of 3:1 (black circle), 3:2 (gray triangle) and 1:1 (red square). **c)** Storage (G’) and loss modulus (G’’) at different time points of hydrogels formation by mixing **1** and **2** at three different BCN/N_3_ ratios (1:1 red square, 3:2 grey triangle, 3:1 black circle) and exposure of the gel to UV light (365nm, 230 mW/cm^2^) for 2 min starting from min 5 until min 7.5 (purple window). **d)** Ratios of storage modulus (G’) and the storage modulus at t=0 (G_0_’) during UV-light (365 nm, 20 mW/cm^2^) exposure with different ration of BCN/N_3_ (1:1 red square, 3:2 grey triangle, 3:1 black circle).

Next, the photodegradation of the hydrogels was studied using a UV-vis lamp with 320-500 nm filter. The light intensity at the end of the optic fiber used during the experiment was set at 20 mW/cm^2^. Using a continuous light exposure of 2 min., a rapid decrease of the storage modulus was observed (Fig. 2c) and had turned into a yellow liquid. The ratio between G’(t) and G’(t_0_) revealed that the degradation does not depend on crosslinker density, achieving complete degradation for all ratios within 2 min. (Fig. 2d).

### Embedding, Photodegration and Retrieval of hMSCs from the Hydrogel

Next, attention was focused on embedding cells in the hydrogel followed by culturing, retrieval and determination of cell viability. Due to the chemical design of the hydrogel platform, gelation only takes place upon addition of crosslinker **2**, making embedding of cells in the 3D biomaterial straightforward. Thus, hMSCs were suspended in a 5% w/v solution of HA-BCN (**1**) in PBS buffer. The relatively low viscosity of the solution, due to the low molecular weight of the employed hyaluronic acid (60 kDa), allowed gentle resuspension of the cell pellet. The resuspended hMSCs were then placed in molds and 0.6 eq of **2** was added which within 5-10 min resulted in gel formation when incubated at 37 °C. Fresh MSC media was added, and the cells were cultured for 24 h. Live and dead staining showed a viability of 87% after 24h in the gel (Fig. 3a), demonstrating cytocompatibility of the hydrogel. To harvest the cells, the hydrogel was exposed to UV light at 30 s intervals at 20 mW/cm^2^. After each time interval, the released cells were collected in fresh media. Approx. 2 to 4 min was needed for complete photodegradation of the hydrogel which is consistent with the rheological data. Once the cells were harvested, they were seeded in a 24 well plate and cultured for up to 72 h (Fig. 3c). After harvesting and reseeding (t = 0), the cells still showed a round phenotype. Nevertheless, after 24 h, the cells attached to the bottom of wells and an elongated morphology was observed. Live and dead staining after 24 h revealed a viability of greater than 78% (Fig. 3b and Fig. S3).

**Figure 3.**
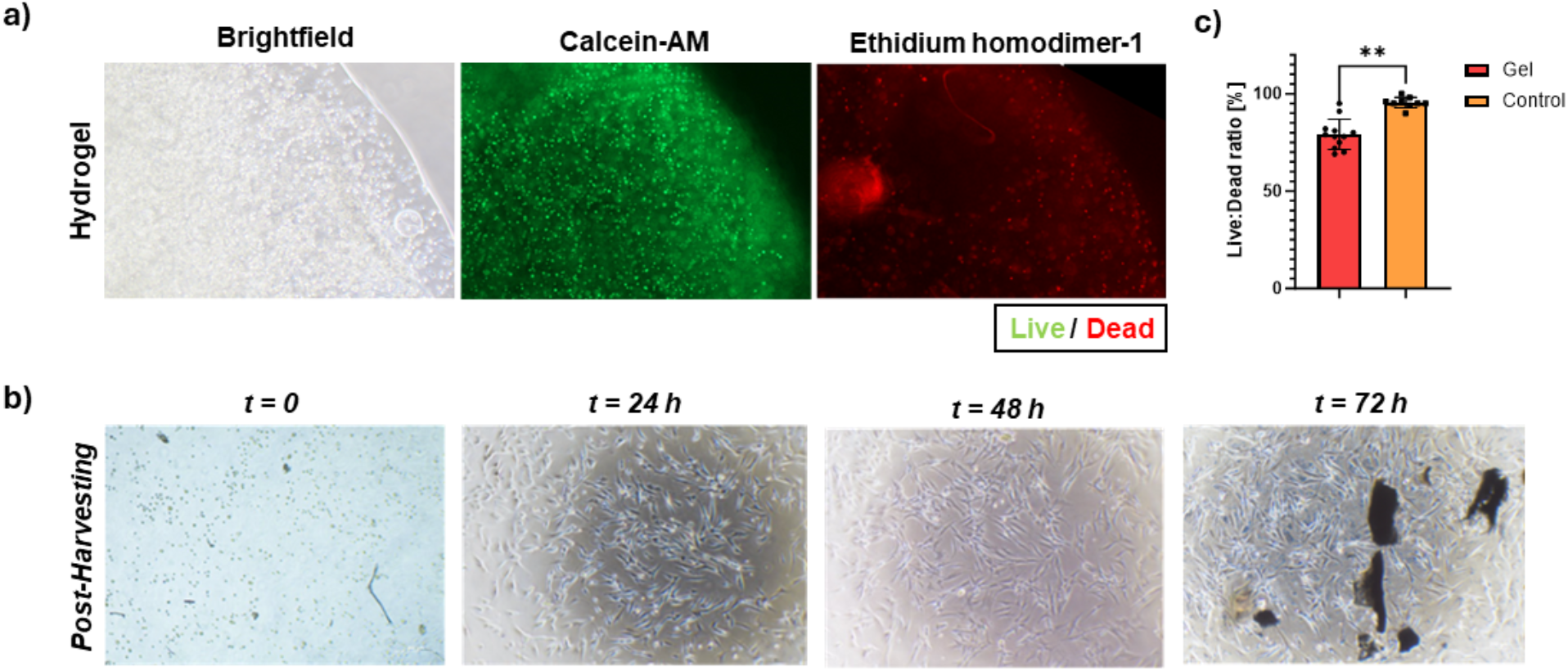
Evaluation of MSC viability and morphology after retrieval from Hydrogel. ***a)*** Brightfield (left) and fluorescence (right) images of MSC after 24h culturing, indicating alive (calcein-AM stained cells in green) and dead cells (ethidium homodimer-1 stained cells in red). **b)** Bar plots representing Live:Dead ratio of 24 h cultured MSCs after being retrieved from hydrogel (Gel) compared to not embedded MSCs (Control). **c**) Bright field images of re-seeded MSCs at 1, 24, 48, and 72 h after retrieved from photocleavable hydrogel.

### Photo-Mediated Softening of Hydrogels using 405 nm Visible Light

Next, we evaluated cell morphology changes induced by photo-mediated softening of the HA-BCN based hydrogel using 405 nm visible light (Fig. S4). First, hydrogel biocompatibility was determined at days 1, 7, and 14. The impact of ECFCs on MSC viability and metabolic activity within the HA-BCN hydrogel was quantified. Thus, cells were gently resuspended at 5×10^6^ cells/mL containing 5% w/v solution of HA-BCN (**1**). In the co-culture, the ratio of ECFCs to MSCs was 1:3 and total cell concentration was kept identical in the monoculture (*i*.*e* 5×10^6^ MSCs/mL). After placing the suspension in 50 μL disc-shaped molds, photosensitive crosslinker **2** and integrin derived peptide **3** were added and the resulting suspension was incubated at 37 ºC, leading to gel formation within 15 min. Cell-laden hydrogels were kept in 24-well plates for 14 days at 37 ºC in EC media for the co-cultures and MSC media for the monocultures (MSCs alone).

The metabolic activity increased over 14 days for the co-cultures, while the trend was not significant in monoculture conditions (Fig. 4a). These finding suggest that cells in the hydrogel are metabolically active and a co-culture with ECFCs enhances this effect, likely due to direct cell-cell interaction and paracrine signaling between both cell types.^3^ Furthermore, live/dead staining revealed an increasing trend in cell viability within co-culture gels over time (Fig. 4b), reaching 82% at day 14 (Fig. 4c), indicating that HA-BCN supports cell viability.

**Figure 4.**
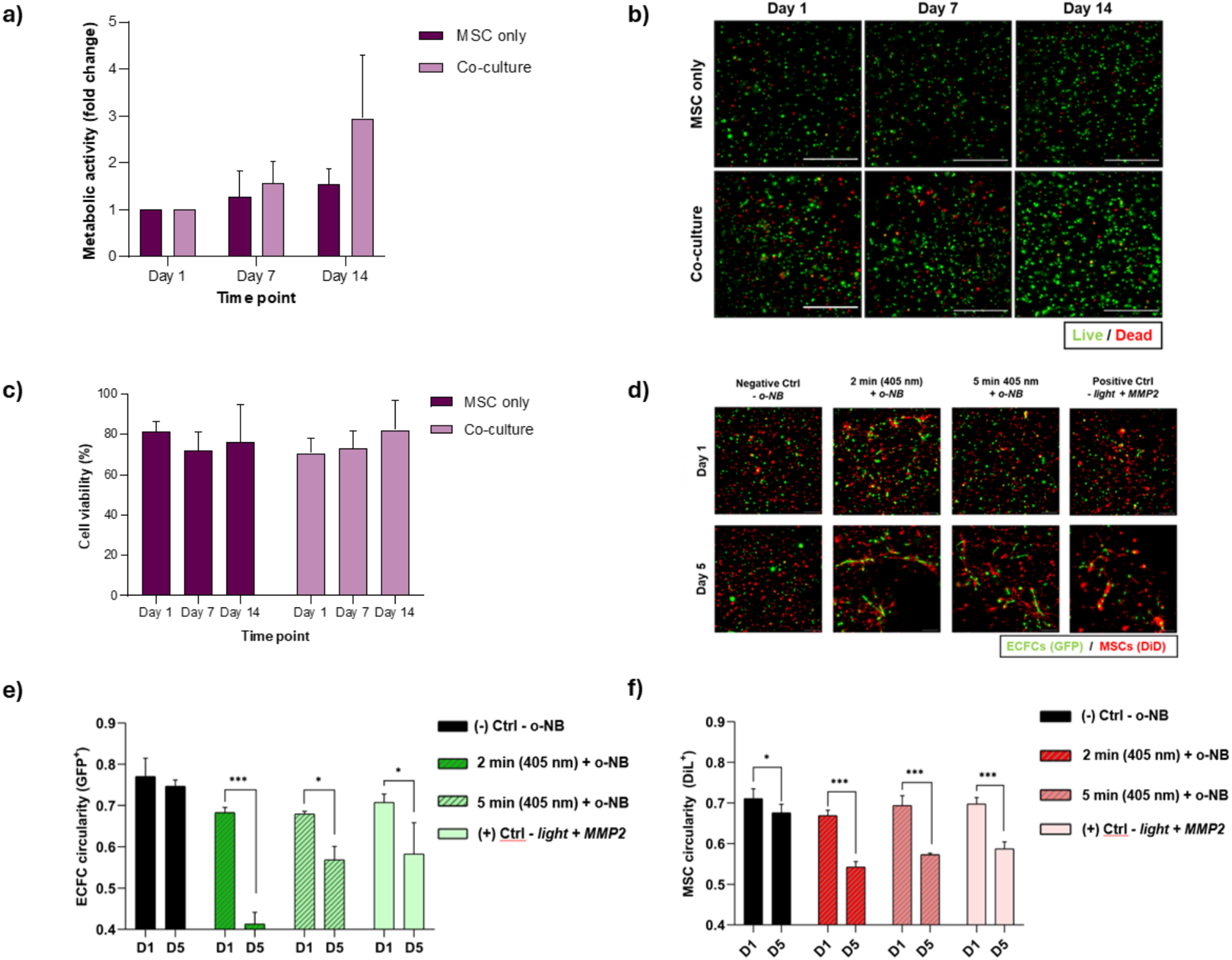
Effect of photo-mediated HA-BCN softening on ECFC and MSC morphology. **(a)** Quantitative analysis of cell metabolic activity within 5% w/v HA-BCN, between monoculture and co-culture conditions (left) and over 14 days (right). Mean fluorescence values ± SD per experimental group expressed as fold change relative to Day 1 are shown. Statistical differences were determined by performing a two-way ANOVA test followed by Sidak’s. **(b)** Z-stack projections of cells within HA-BCN gels after live/dead staining at days 1, 7, and 14. Live cells were labeled with Calcein (green) and dead cells with Ethidium Homodimer-1 (red). Scale bar=500 μm. **(c)** Quantitative analysis of cell viability within 5% w/v HA-BCN, between monoculture and co-culture conditions (top) and over 14 days (bottom). Mean viability values ± SD per experimental group is shown. Statistical differences were determined by performing a two-way ANOVA test followed by Sidak’s (top) and Tukey’s (bottom) multiple comparison tests. **(d)** Z-stack projections of cells within HA-BCN gels at days 1 and 5 after inducing photo-mediated softening for 2 and 5 min. ECFCs express GFP (green) and MSCs are stained with DiL (red). Scale bar=200 μm. **(e)** and **(f)** Quantitative analyses of ECFC and MSC circularity, respectively, over 5 days within 3.8% w/v HA-BCN gels after inducing photo-mediated softening for 2 and 5 min. Mean circularity values ± SD per experimental group calculated taking the inverse of the average cell aspect ratio are shown. Statistical differences were determined by performing a two-way ANOVA test (*p<0.05, *** p<0.001).

Next, we focused on inducing photo-mediated softening in HA-BCN hydrogels to determine morphological changes in cells over time. Compounds **2** and **3** were mixed with 3.8% w/v HA-BCN to form photodegradable gels, while Polyoxyethylene (2kDa) bis(azide) (N_3_-PEG-N_3_) was used instead of compound **2** as a negative control to form non-photosensitive hydrogels. For positive control gels, we used a 50% molar ratio of compound **2** and **4**. After crosslinking the samples for 10 min at 37 ºC, the photodegradable samples were exposed to light (λ = 405 nm, I = 140 μJs^-1^cm^-2^) for 2 and 5 min. All samples were then incubated for 5 days at 37 ºC in 2% FBS EGM-2 medium and cell circularity was quantified at days 1 and 5.

ECFC circularity significantly decreased at day 5 in photodegradable and positive control samples (Fig. 4e), while cells within negative control samples still exhibited a round phenotype (Fig. 4d), indicating that a reduction in gel stiffness induces ECFC stretching. MSC circularity significantly decreased at day 5 in all conditions, with the greatest reduction observed in photodegradable and positive control samples (Fig. 4f). These results indicate that hydrogel stiffness influences the morphology of MSC and ECFC 3D co-culture, with the potential to guide cell behaviour through photo-mediated changes in stiffness.^1^

## CONCLUSIONS

In this study, we synthesized hyaluronic acid (HA) derivatized with BCN (**1**) and a photosensitive crosslinker (**2**) having azides that can form hydrogels with tuneable mechanical properties. The hydrogels exhibit fast photodegradation upon exposure to UV light (365 nm), achieving complete breakdown within minutes regardless of crosslinker density. This feature enabled the efficient retrieval of human mesenchymal stromal cells (hMSCs) embedded within the hydrogels, maintaining high cell viability post-retrieval. This property is essential for a hydrogel platform aimed at employing 3D biological experiments, allowing for a rapid harvesting of cultured cell with minimal perturbation of cell properties. The cytocompatibility of the hydrogels was further confirmed through long-term culture experiments, showing significant cell proliferation and viability, especially in a co-culture system involving endothelial colony-forming cells (ECFCs) and MSCs. Additionally, photo-mediated softening of the hydrogels led to notable changes in cell morphology, highlighting the potential to guide cellular behaviour through visible light (405 nm) mediated mechanical modulation. The incorporation of the RGD cell adhesive peptide in these experiments excluded the possibility that the change in morphology is only due to adhesive properties. It shows that the softening of the material is essential to promote the phenotypical differentiation, which can be achieved also by endogenous enzymes produced by cells when MMP2 cleavable peptide was used as a secondary crosslinker in a positive control experiment. The finding will open new venues and for example a range of different peptides or proteins can be incorporated to direct cell attachment thereby controlling differentiation. Fine-tuning the photodegradation kinetics and mechanical properties could lead to more sophisticated control over the cell microenvironment, benefiting tissue engineering and regenerative medicine applications. Moreover, the development of *in vivo* models to assess the performance and biocompatibility of these hydrogels could pave the way for clinical applications in wound healing, drug delivery, and cell therapy.

## ASSOCIATED CONTENT

### Supporting Information

The Supporting Information is available with:

Supplementary figures, reaction schemes, chemical procedures, and copies of NMR spectra (PDF).

## AUTHOR INFORMATION

## Author Contributions

F.P., P.C., T.V., R.L., and G.J.B. designed the project. G.J.B. was responsible for overall project management. F.P. performed the synthesis and the characterization of the molecules. F.P. and P.C. performed the mechanical characterization of the material. F.P., S.B., and S.M. performed the cell harvesting experiments. T.V. and R.L. designed the mechanical tests and the hydrogel softening experiments. M.G. and L.V.d.R. performed the softening of the cocultured cells. The manuscript was written by F.P., S.B., L.V.d.R., and G.J.B. All authors reviewed the final manuscript.

## Notes

The authors declare no competing financial interest.

## ACKNOWLEDGMENT

This work was supported by the EC H2020 Marie Curie doctorate COFUND RESCUE programme ‘Regenerating Medicine Training Network’.

## REFERENCES

(1) Lee, K. Y.; Mooney, D. J. Hydrogels for tissue engineering. Chem. Rev. 2001, 101 (7), 1869–1879.

(2) Drury, J. L.; Mooney, D. J. Hydrogels for tissue engineering: Scaffold design variables and applications. Biomaterials 2003, 24 (24), 4337–4351.

(3) Tibbitt, M. W.; Anseth, K. S. Hydrogels as extracellular matrix mimics for 3D cell culture. Biotechnol. Bioeng. 2009, 103 (4), 655–663.

(4) Annabi, N.; Tamayol, A.; Uquillas, J. A.; Akbari, M.; Bertassoni, L. E.; Cha, C.; Camci-Unal, G.; Dokmeci, M. R.; Peppas, N. A.; Khademhosseini, A. 25th anniversary article: Rational design and applications of hydrogels in regenerative medicine. Adv. Mater. 2014, 26 (1), 85–123.

(5) Khademhosseini, A.; Langer, R. A decade of progress in tissue engineering. Nat. Protoc. 2016, 11 (10), 1775–1781.

(6) Revete, A.; Aparicio, A.; Cisterna, B. A.; Revete, J.; Luis, L.; Ibarra, E.; Segura Gonzalez, E. A.; Molino, J.; Reginensi, D. Advancements in the use of hydrogels for regenerative medicine: Properties and biomedical applications. Int. J. Biomater. 2022, 2022, 3606765.

(7) Chai, Q.; Jiao, Y.; Yu, X. Hydrogels for biomedical applications: Their characteristics and the mechanisms behind them. Gels 2017, 3 (1), 6.

(8) Das, S.; Kumar, V.; Tiwari, R.; Singh, L.; Singh, S. Recent advances in hydrogels for biomedical applications. Asian J. Pharm. Clin. Res. 2018, 11 (11), 62.

(9) Mandal, A.; Clegg, J. R.; Anselmo, A. C.; Mitragotri, S. Hydrogels in the clinic. Bioeng. Transl. Med. 2020, 5 (2), e10158.

(10) Vermonden, T.; M, N. A.; van, M. J.; Hennink, W. E. Rheological studies of thermosensitive triblock copolymer hydrogels. Langmuir 2006, 22 (24), 10180–10184.

(11) Fan, X.; Zhu, L.; Wang, K.; Wang, B.; Wu, Y.; Xie, W.; Huang, C.; Chan, B. P.; Du, Y. Stiffness-controlled thermoresponsive hydrogels for cell harvesting with sustained mechanical memory. Adv. Healthc. Mater. 2017, 6 (5), 1601152.

(12) Klouda, L. Thermoresponsive hydrogels in biomedical applications: A seven-year update. Eur. J. Pharm. Biopharm. 2015, 97 (Pt B), 338–349.

(13) Arrigo, A. P.; Suhan, J. P.; Welch, W. J. Dynamic changes in the structure and intracellular locale of the mammalian low-molecular-weight heat shock protein. Mol. Cell. Biol. 1988, 8 (12), 5059–5071.

(14) Al-Fageeh, M. B.; Marchant, R. J.; Carden, M. J.; Smales, C. M. The cold-shock response in cultured mammalian cells: harnessing the response for the improvement of recombinant protein production. Biotechnol. Bioeng. 2006, 93 (5), 829–835.

(15) Huang, J.; Jiang, X. Injectable and degradable pH-responsive hydrogels via spontaneous amino-yne click reaction. ACS Appl. Mater. Interfaces 2018, 10 (1), 361–370.

(16) Shi, Y. G.; Li, D.; Ding, J. F.; He, C. L.; Chen, X. S. Physiologically relevant pH- and temperature-responsive polypeptide hydrogels with adhesive properties. Pol. Chem. 2021, 12 (19), 2832–2839.

(17) Kruse, C. R.; Singh, M.; Targosinski, S.; Sinha, I.; Sorensen, J. A.; Eriksson, E.; Nuutila, K. The effect of pH on cell viability, cell migration, cell proliferation, wound closure, and wound reepithelialization: In vitro and in vivo study. Wound Repair Regen. 2017, 25 (2), 260–269.

(18) Beninatto, R.; Barbera, C.; De Lucchi, O.; Borsato, G.; Serena, E.; Guarise, C.; Pavan, M.; Luni, C.; Martewicz, S.; Galesso, D.; Elvassore, N. Photocrosslinked hydrogels from coumarin derivatives of hyaluronic acid for tissue engineering applications. Mater. Sci. Eng. C Mater. Biol. Appl. 2019, 96, 625–634.

(19) Rapp, T. L.; DeForest, C. A. Tricolor visible wavelength-selective photodegradable hydrogel biomaterials. Nat. Commun. 2023, 14 (1), 5250.

(20) Thomas, S. W. New Applications of Photolabile Nitrobenzyl Groups in Polymers. Macromol. Chem. Phys. 2012, 213 (23), 2443–2449.

(21) Zhao, H.; Sterner, E. S.; Coughlin, E. B.; Theato, P. o-nitrobenzyl alcohol derivatives: Opportunities in polymer and materials science. Macromolecules 2012, 45 (4), 1723–1736.

(22) Claassen, C.; Claassen, M. H.; Gohl, F.; Tovar, G. E. M.; Borchers, K.; Southan, A. Photoinduced cleavage and hydrolysis of o-nitrobenzyl linker and covalent linker immobilization in gelatin methacryloyl hydrogels. Macromol. Biosci. 2018, 18 (9), e1800104.

(23) Lunzer, M.; Shi, L.; Andriotis, O. G.; Gruber, P.; Markovic, M.; Thurner, P. J.; Ossipov, D.; Liska, R.; Ovsianikov, A. A modular approach to sensitized two-photon patterning of photodegradable hydrogels. Angew. Chem. Int. Ed. 2018, 57 (46), 15122–15127.

(24) Hansen, M. J.; Velema, W. A.; Lerch, M. M.; Szymanski, W.; Feringa, B. L. Wavelength-selective cleavage of photoprotecting groups: strategies and applications in dynamic systems. Chem. Soc. Rev. 2015, 44 (11), 3358–3377.

(25) Cadet, J.; Sage, E.; Douki, T. Ultraviolet radiation-mediated damage to cellular DNA. Mutat. Res. 2005, 571 (1-2), 3–17.

(26) D’Orazio, J.; Jarrett, S.; Amaro-Ortiz, A.; Scott, T. UV radiation and the skin. Int. J. Mol. Sci. 2013, 14 (6), 12222–12248.

(27) Wang, P. W.; Hung, Y. C.; Lin, T. Y.; Fang, J. Y.; Yang, P. M.; Chen, M. H.; Pan, T. L. Comparison of the biological impact of UVA and UVB upon the skin with functional proteomics and immunohistochemistry. Antioxidants 2019, 8 (12), 569.

(28) Burdick, J. A.; Prestwich, G. D. Hyaluronic acid hydrogels for biomedical applications. Adv. Mater. 2011, 23 (12), H41–H56.

(29) Highley, C. B.; Prestwich, G. D.; Burdick, J. A. Recent advances in hyaluronic acid hydrogels for biomedical applications. Curr. Opin. Biotechnol. 2016, 40, 35–40.

(30) Trombino, S.; Servidio, C.; Curcio, F.; Cassano, R. Strategies for hyaluronic acid-based hydrogel design in drug delivery. Pharmaceutics 2019, 11 (8), 407.

(31) Dommerholt, J.; Schmidt, S.; Temming, R.; Hendriks, L. J.; Rutjes, F. P.; van Hest, J. C.; Lefeber, D. J.; Friedl, P.; van Delft, F. L. Readily accessible bicyclononynes for bioorthogonal labeling and three-dimensional imaging of living cells. Angew. Chem. Int. Ed. 2010, 49 (49), 9422–9425.

(32) LeValley, P. J.; Neelarapu, R.; Sutherland, B. P.; Dasgupta, S.; Kloxin, C. J.; Kloxin, A. M. Photolabile linkers: Exploiting labile bond chemistry to control mode and rate of hydrogel degradation and protein release. J. Am. Chem. Soc. 2020, 142 (10), 4671–4679.

(33) Cabral-Pacheco, G. A.; Garza-Veloz, I.; Castruita-De la Rosa, C.; Ramirez-Acuna, J. M.; Perez-Romero, B. A.; Guerrero-Rodriguez, J. F.; Martinez-Avila, N.; Martinez-Fierro, M. L. The roles of matrix metalloproteinases and their inhibitors in human diseases. Int. J. Mol. Sci. 2020, 21 (24), 9739.

(34) Lv, H.; Wang, H.; Zhang, Z.; Yang, W.; Liu, W.; Li, Y.; Li, L. Biomaterial stiffness determines stem cell fate. Life Sci. 2017, 178, 42–48.

(35) Ma, Y.; Lin, M.; Huang, G.; Li, Y.; Wang, S.; Bai, G.; Lu, T. J.; Xu, F. 3D spatiotemporal mechanical microenvironment: A hydrogel-based platform for guiding stem cell fate. Adv. Mater. 2018, 30 (49), e1705911.

(36) Chang, C. Y.; Lin, C. C. Hydrogel models with stiffness gradients for interrogating pancreatic cancer cell fate. Bioengineering 2021, 8 (3), 37.

