## Supplementary material for "A Light Sensitive Hyaluronic Acid-based Hydrogel for 3D Culturing, Differentiation, and Harvesting of Mesenchymal Stromal Cells": SI

**Table of contents** page


**1) Supplementary Figures** S3

Figure S1. Hydrogel photodegradation using UV light S3

Figure S2. Frequency sweep and oscillation sweep of the hydrogels S3

Figure S3. Life and death assay of cells after the hydrogel degradation S4

Figure S4. Compression modulus measured via Dynamic Mechanical Analysis S4

**2) Chemical Synthesis** S5

Scheme S1. Synthesis of BCN-$NH_2$ (**4**) ` S5

**3) Synthesis of Photocleavable Crosslinker (PC-CL) (2)** S8

Scheme S2. Synthesis of PC-CL (**2**) S8

**4) SPPS of Peptides 3 ($N_3$-GGRGDSP) and 4 (AcK($N_3$)RDIPVSLRSGDRK($N_3$))** S12

Scheme S3. General scheme for the SPPS of compounds **3** and **4** S12

**5) References** S14

**6) NMR Spectra** S15

### 1) Supplementary Figures

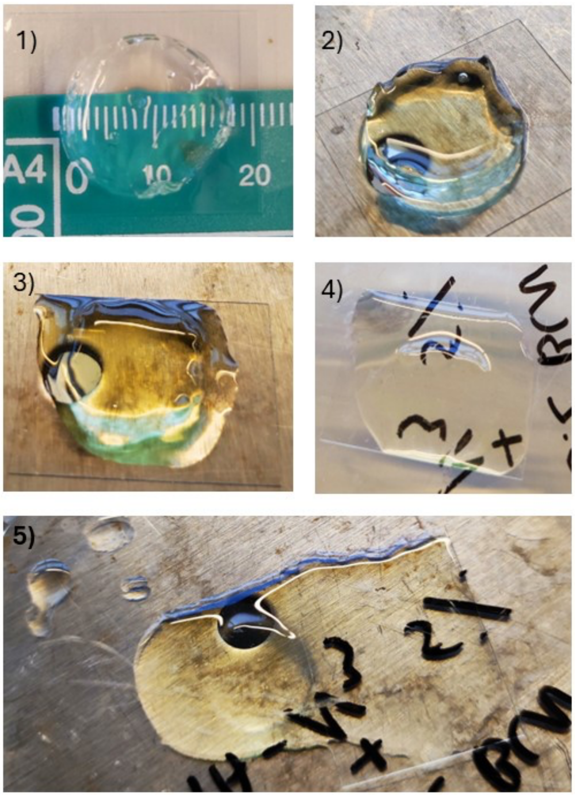


**Figure S1.** Hydrogel photodegradation using UV light (365 nm 20 mW/cm$^2$) after 30 sec (1) 60 sec (2) 90 sec (3) 120 sec (4) 150 sec (5).

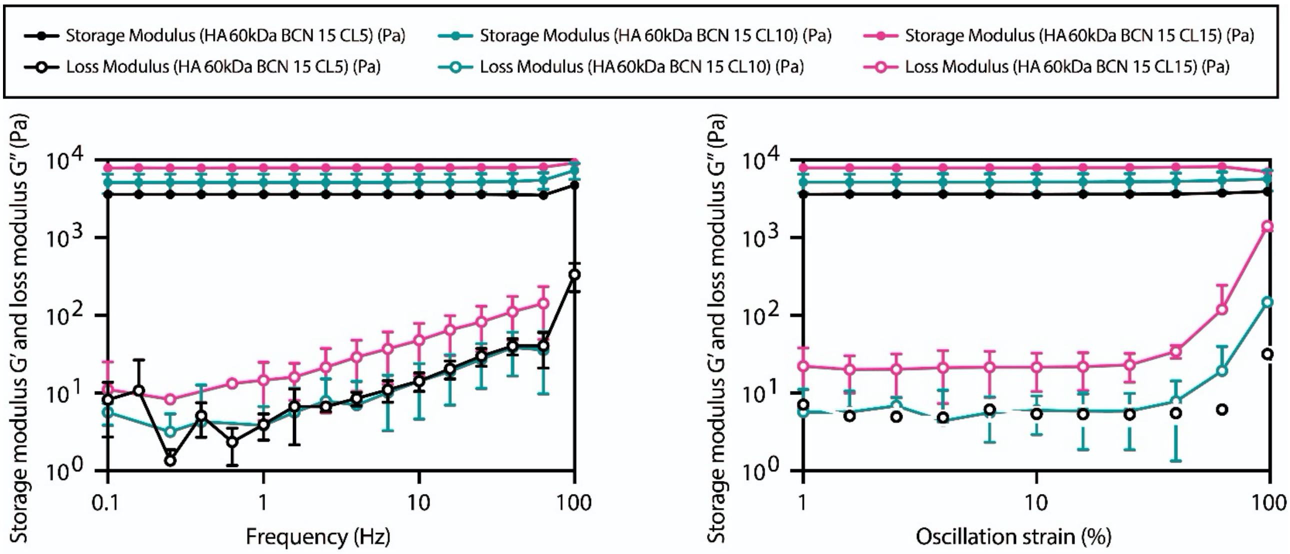


**Figure S2.** Frequency sweep (left) and oscillation sweep (right) of the hydrogels made by mixing HA-BCN (compound **1**) and PC-CL (compound **2**) in a ratio of 1:1 (pink triangle) 3:2 (green circle) and 3:1 (black circle).

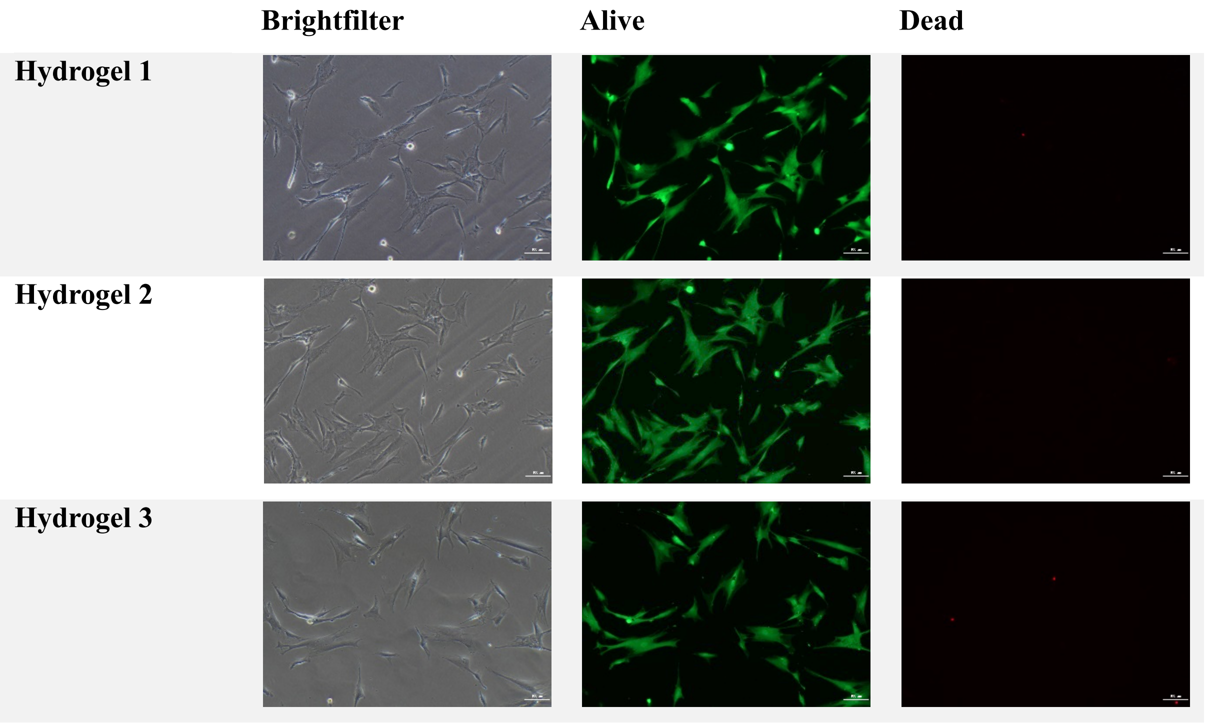


**Figure S3.** Life and death assay of cells retrieved from three different samples after the hydrogel degradation using UV 365nm (20 mW/cm$^2$).

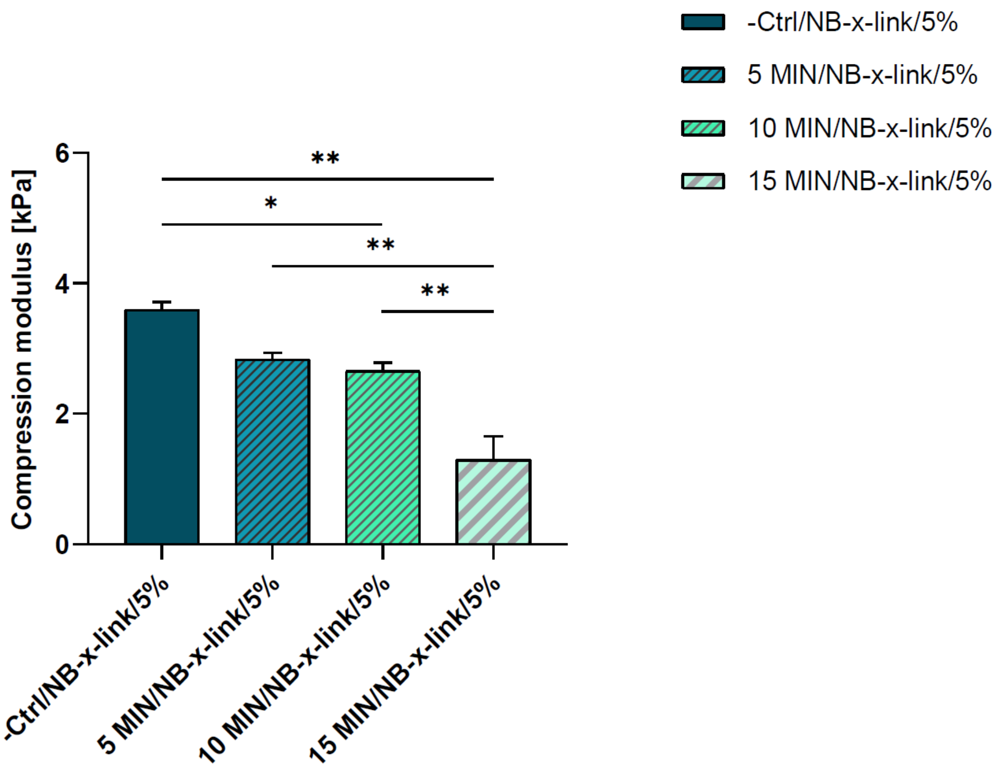


**Figure S4.** Compression modulus measured via Dynamic Mechanical Analysis (DMA) after 5, 10, and 15 min of 405 nm light (140 µJ*s-1*cm-2) exposition in a light box of hydrogels constituted by mixing compounds **1** and **2** in a ratio of 3:1.

### 2) Chemical Synthesis

**General Methods and Materials.** All reagents and solvents were purchased from commercial sources and used without further purification. Anhydrous reactions were carried out under an argon atmosphere. Reactions were monitored using TLC on aluminium-backed plates coated with Silica Gel 60 F254 (E. Merck) and visualized by UV light (254 nm) where applicable, and by application of 5% $H_2SO_4$ in EtOH or with a solution of $(NH_4)_6Mo_7O_{24}.H_2O$ (25.0 $gL^{-1}$) in 10% $H_2SO_4$ in EtOH, as appropriate, and heating. Column chromatography was performed on silica gel G60 (Silicycle, 60-200 μm, 60 Å), or on Bondapak C-18 (Waters). Size exclusion chromatography was carried out on a Sephadex™ LH-20 using MeOH/DCM (1/1, v/v) as elution system. NMR spectra were collected on an Agilent 400-MR, Varian or Bruker 600 UltraShield. Signals are reported in terms of chemical shift [δ in parts per million (ppm)] relative to tetramethylsilane (TMS) as the internal standard. NMR data are presented as follows: Chemical shift, multiplicity (s = singlet, br. s = broad singlet, d = doublet, t = triplet, dd = doublet of doublet, m = multiplet and/or multiple resonances), coupling constant in hertz (Hz), integration. All NMR signals were assigned on the basis of $^1H$ NMR, $^{13}C$ NMR, COSY, TOCSY and HSQC experiments. Mass spectra were recorded on an ABISciex 5800 MALDI-TOF-TOF or Shimadzu LCMS-IT-TOF mass spectrometer. The matrix used for MALDI-TOF MS was 2,5-dihydroxy-benzoic acid (DHB) and ultramark 1621 as the internal standard.

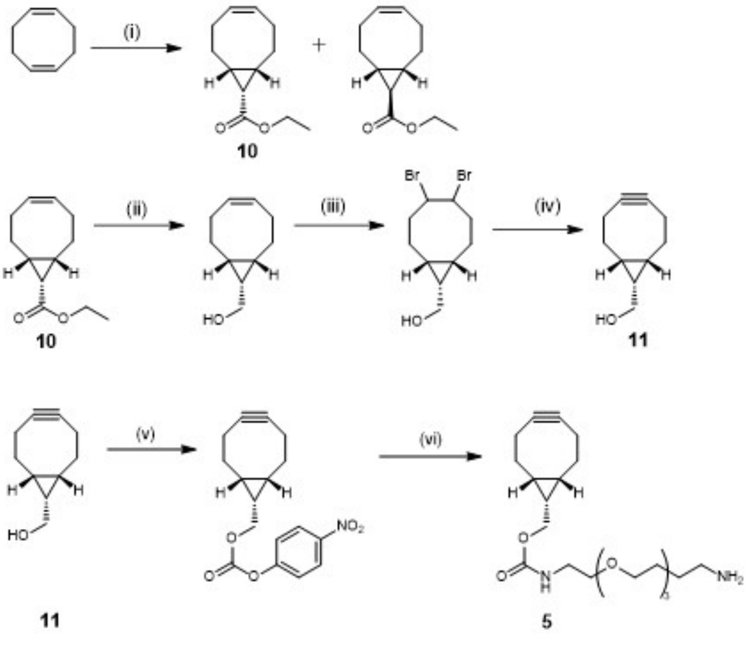


**Scheme S1.** Synthesis of BCN-$NH_2$ (**4**). Reagents and conditions: (i) Cyclooctadiene, $Rh(II)_2(OAc)_4$, DCM, 0 °C to RT, 72 h. (ii) $LiAlH_4$, $Et_2O$, 0 °C, 15 min. (iii) $Br_2$ , DCM, 0 °C, 15 min (iv) KOtBu, THF, reflux, 2 h. (v) 4-Nitrophenyl chloroformate, pyridine, DCM, 0 °C to RT, 2 h. (vi) 1,8-diamino-3,6-dioxaoctane, DMF.

**Endo-BCN-OH Synthesis (11).** (1R,8S,9s)-Bicyclo[6.1.0]non-4-yn-9-ylmethanol (Endo-BCN-OH) was synthesized as previously described.[1] Ethyl diazoacetate (5.81 g, 51 mmol, 5.4 mL) was dissolved in DCM (30 mL) and added dropwise under inert atmosphere at 0 ºC to a dry 500 mL round bottom flask containing rhodium (II) acetate dimer (1 g, 2.3 mmol) dissolved in 1,5-cyclooctadiene (44 g, 408 mmol, 50 mL). The reaction was stirred at room temperature (RT) for 72 h. Subsequently, the reaction mixture was filtered through silica and concentrated under reduced pressure. The product was purified by column chromatography using toluene, resulting in two stereoisomers: (**10**) endo (2.9 g, 14.9 mmol, 29% yield) and exo (3.5 g, 18 mmol, 35% yield). (**1**): $^1$H NMR (400 MHz, $CDCl_3$): δ 5.60 – 5. 56 (m, 2H), 4.14 – 4.07 (q, 2H), 2.55 – 2.44 (m, 2H), 2.25 - 2.14 (m, 2H), 2.09 – 1.99 (m, 2H), 1.87 – 1.76 (m, 2H), 1.72 – 1.66 (t, 1H), 1.43 – 1.32 (m, 2H), 1.28 – 1.22 (t, 3H). $^{13}$C NMR (75 MHz, $CDCl_3$): δ 172.2, 129.4, 59.6, 27.0, 24.1, 22.6, 21.2, 14.4 (**2**): $^1$H NMR (400 MHz, $CDCl_3$): δ 5.65 – 5.53 (m, 2H), 4.14 – 4.04 (q, 2H), 2.54 – 2.43 (m, 2H), 2.22 - 2.13 (m, 2H), 2.12 – 2.02 (m, 2H), 1.60 – 1.52 (m, 2H), 1.51 – 1.39 (m, 2H), 1.27 – 1.20 (t, 3H), 1.18 – 1.13 (t, 1H). $^{13}$C NMR (75 MHz, $CDCl_3$): δ 174.3, 129.8, 60.1, 28.2, 27.8, 27.6, 26.6, 14.2. $LiAlH_4$ (582 mg, 15.35 mmol) was dissolved in anhydrous diethyl ether (50 mL) and then added dropwise at 0 ºC to a solution of endo-intermediate (**1**) (2.9 g, 14.9 mmol) in anhydrous diethyl ether (50 mL). The reaction was stirred at RT for 30 min. and then cooled to 0 ºC with simultaneous addition of water (4 mL) during which the grey solid turned white. The mixture was dried ($Na_2SO_4$), filtered, and the filtrate was concentrated under reduced pressure to give compound **11** as yellow oil. The crude material was dissolved in DCM (100 mL) and a solution of $Br_2$ (0.9 mL, 18 mmol) in DCM (20 mL) was added dropwise at 0 ºC. The reaction mixture gradually turned red upon the addition. The reaction was quenched with 10% aqueous solution of $Na_2S_2O_3$ (30 mL) after TLC analysis showed compete consumption of the starting material. The solution was extracted with DCM (3 x 60 mL) and the combined organic layers were dried ($Na_2SO_4$), filtered and the filtrate was concentrated under reduced pressure to give **3** in white amorphous crystals. Without further purification, the material was dissolved in THF (120 mL) and placed under inert atmosphere of nitrogen. Subsequently, 1 M solution of KOtBu (40.5 mmol, 40.5 mL) was added dropwise at 0 ºC. The reaction mixture was stirred under reflux for 2 h, after which it was quenched with a saturated solution of aqueous $NH_4Cl$. The mixture was extracted with DCM (3 x 50 mL) and the combined organic layers were dried ($MgSO_4$) and concentrated under reduced pressure. The residue was purified by silica gel column chromatography (eluent: petroleum ether:EtOAc), yielding compound **11** as amorphous crystals (0.82 g, 5.46 mmol, 37% yield over 3 steps). $^1$H NMR (400 MHz, $CDCl_3$): δ 3.73 (d, J=7.9 Hz, 2H), 2.37 – 2.15 (m, 4H), 1.68

– 1.51 (m, 3H), 1.40 –1.27 (m, 2H), 1.18 (bs, 1H), 0.99-0.89 (m, 2H). $^{13}$C NMR ($CDCl_3$, 75MHz): δ 99.03, 60.16, 29.2, 21.65, 21.56, 20.18.

**Endo-BCN-PEG$_2$-NH$_2$ (5).**[1] Compound **11** (0.82 g, 5.46 mmol) was dissolved in DCM (15 mL) and pyridine (0.98 mL, 1.36 mmol) and *p*-nitrophenyl chloroformate (1.42 g, 7.1 mmol) was added. The reaction mixture was stirred at RT for 30 min, after which it was quenched with a saturated solution of $NH_4Cl$. The mixture was extracted with DCM (3 x 20 mL) and combined organic layers were dried ($MgSO_4$) and concentrated under reduced pressure. Without further purification, the material was dissolved in DMF (10 mL) and added dropwise to a solution of 1,8-diamino-3,6-dioxaoctane (3.83 g, 25.86 mmol) and triethylamine (1.82 mL,12.9 mol) in DMF (10 mL) and left stirring at RT for 30 min. The mixture was concentrated under reduced pressure, after which it was taken up in a minimal amount of $CH_2Cl_2$ (100 mL) and extracted with 1M aqueous solution of NaOH (2 x 200 mL) and water (1 x 200 mL). The combined organic phases were dried ($MgSO_4$) and concentrated under reduced pressure. The residue was purified with silica gel column chromatography (Eluent: DCM:MeOH; 1% $Et_3N$), yielding the final product N-[(1R,8S,9S)-Bicyclo[6.1.0]non-4-yn-9-ylmethyloxycarbonyl]-1,8-diamino-3,6-dioxaoctane (**5**) referred to as BCN-$NH_2$, as a light yellow oil (1.24 g, 0.4 mmol, 74% yield over two steps). $^1$H NMR (600 MHz, $CDCl_3$) δ 5.41 (bs, 1H) 4.13 (d, J = 8.1 Hz, 2H), 3.64 – 3.58 (m, 4H), 3.55 (t, J = 5.1 Hz, 2H), 3.52 (t, J = 5.1 Hz, 2H), 3.37 (q, J = 5.4 Hz, 2H), 2.88 (t, J = 5.1 Hz, 2H), 2.32 – 2.16 (m, 6H), 1.62 – 1.52 (m, 2H), 1.34 (m, 1H), 0.93 (m, 2H).

#### 3) Synthesis of Photocleavable Crosslinker (PC-CL) (2)

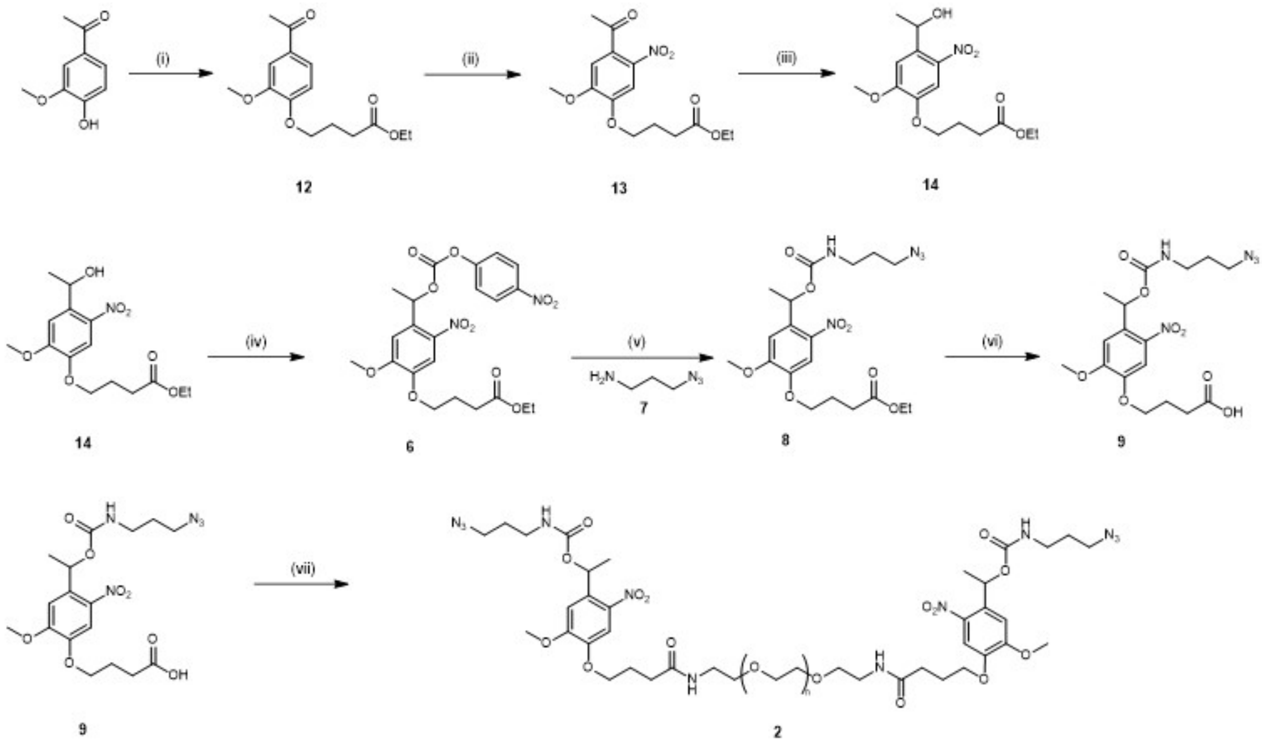

**Scheme S2.** Synthesis of PC-CL (**2**). Reagents and conditions: (i) Acetovanillone, Ethyl 4-bromobutyrate, $K_2CO_3$, DMF, R.T. Overnight (ii) TFA, $NaNO_3$ (iii) DCM/MeOH, $NaBH_4$ (iv) 4-nitrophenyl chloroformate, DCM, TEA (v) compound **7** (2eq.) (vi) LiOH, THF, $H_2O$ (vii) DMF, DIPEA, HATU, HOAt, $NH_2$-PEG(1kDa)-$NH_2$.

**Synthesis of Compound 12.**[2] Acetovanillone (10 g, 60 mmol) was dissolved in of anhydrous DMF (50 mL). $K_2CO_3$ (12.5g, 72 mmol) was added and the ethyl 4-bromobutyrate (10.5 mL, 72 mmol) was slowly added and the mixture was left stirring at RT overnight. The reaction mixture was concentrated under reduced pressure, and the residue was diluted with EtOAc (100 mL) and extracted 3 times with water. The organic phase was dried ($MgSO_4$) and filtrated and the filtrate was concentrated under reduced pressure to give white amorphous crystals. The crystals were then triturated using a mortar and washed with water multiple times to obtain compound **12** as white powder (14.6 g, 87%). $^1$H NMR (400 MHz, $CDCl_3$) δ 7.59 – 7.50 (m, 2H), 6.89 (d, J = 8.3 Hz, 1H), 4.18 – 4.11 (m, 4H), 3.91 (s, 3H), 2.57 – 2.51 (t, 2H), 2.56 (s, 3H) 2.18 (p, J = 6.8 Hz, 2H), 1.25 (t, J = 7.1 Hz, 3H). $^{13}$C NMR (101 MHz, $CDCl_3$) δ 196.78, 172.99, 152.62, 149.28, 130.50, 123.17, 111.26, 110.47, 77.19, 67.79, 60.47, 55.98, 30.56, 26.18, 24.28, 14.18.

**Synthesis of Compound 13.**[2] A portion of compound **12** (1.28 g, 4.57 mmol) was slowly added to ice cold TFA (5 mL) and then $NaNO_3$ (1.16 g, 13.6 mmol) was added. The reaction mixture was allowed to reach RT and left stirring until TLC showed full consumption of the starting material. Next, ice-cold water was added to the reaction mixture until precipitation of a fine yellow solid was observed. The solid was filtrated off and washed extensively with water and left in the desiccator overnight to dry to yield compound **13** (1.06 g, 72% yield). $^1H$ NMR (400 MHz, $CDCl_3$) δ 7.60 (s, 1H), 6.73 (s, 1H), 4.21 – 4.10 (m, 4H), 3.99 – 3.96 (rotamers) (s, 3H), 2.53 (t, J = 7.2 Hz, 2H), 2.48 (s, J = 0.6 Hz, 3H), 2.25 – 2.13 (m, 2H), 1.26 (t, J = 7.1 Hz, 3H).

**Synthesis of Compound 14.**[2] Compound **13** (1.06 g, 3.26 mmol) was dissolved in DCM/MeOH (1:1, 15 mL). The reaction mixture was cooled to 0° prior the addition of $NaBH_4$ (250 mg, 6.5 mmol). Once the TLC showed complete consumption of the starting material, the material was diluted in EtOAc (75 mL) and washed two times with $NH_4Cl$ 1 M solution and two times with brine. The organic phase was dried ($MgSO_4$), filtrated, and the filtrate concentrated under reduced pressure. The compound was recrystallized from EtOH by adding water to obtain compound **14** as a yellow solid (1.05 g, 3.26 mmol, 100%). $^1H$ NMR (400 MHz, $CDCl_3$) δ 7.57 (s, 1H), 6.98 (s, 1H), 6.36 (q, J = 6.4 Hz, 1H), 5.30 (d, J = 0.6 Hz, 1H), 4.92 (s, 1H), 4.20 – 4.06 (m, 4H), 3.9 (s, 3H), 3.35 (t, J = 6.5 Hz, 2H), 3.24 (q, J = 5.0, 3.2 Hz, 2H), 2.53 (t, J = 7.2 Hz, 2H), 2.23 – 2.12 (m, 2H), 1.77 (p, J = 6.6 Hz, 2H), 1.59 (d, J = 7.9 Hz, 4H), 1.26 (td, J = 7.2, 0.6 Hz, 3H).

**Synthesis of Compound 6.**[2] Compound **14** (1.05 g, 3.26 mmol) was dissolved in DCM and triethylamine (680 µL, 4.9 mmol) was added to the solution. The reaction mixture was cooled to 0°C and 4-nitrophenyl chloroformate (800 mg, 4 mmol) was added. The reaction mixture was allowed to reach RT and left stirring overnight. The mixture was filtrated, and the filtrate concentrated under reduced pressure and purified via silica column chromatography using a gradient from 0 to 20% EtOAc in toluene yielding compound **6** as light yellow solid (1 g, 2.03 mmol, 62% yield). $^1H$ NMR (600 MHz, $CDCl_3$) δ 8.26 (d, J = 9.0 Hz, 2H), 7.61 (s, 1H), 7.34 (d, J = 9.0 Hz, 2H), 7.10 (s, 1H), 6.54 (q, J = 6.4 Hz, 1H), 4.20 – 4.11 (m, 4H), 3.99 (s, 3H), 2.54 (t, J = 7.2 Hz, 2H), 2.19 (m, 2H), 1.77 (d, J = 6.4 Hz, 3H), 1.27 (t, J = 7.1 Hz, 3H). $^{13}C$ NMR (600 MHz, $CDCl_3$) δ 125.34, 121.66, 108.99, 108.00, 73.80, 68.35, 60.61, 56.53, 30.58, 24.25, 22.03, 14.24.

**Synthesis of Compound 7.**[2] 3-Bromo propylamine hydrobromide (3.28 g, 15 mmol) was diluted in deionized water (10 mL). $NaN_3$ (3.25 g, 50 mmol) was dissolved in deionized water (15 mL) and added to the mixture. The reaction mixture was heated under reflux overnight. Next, 1 M aqueous sodium hydroxide solution (25 mL) and the resulting solution was extracted using 3x 50 mL of DCM. The combined organic phases were dried ($MgSO_4$), filtrated and the filtrate concentrated to obtain compound **7** as a colorless oil (1.5 g, 15 mmol, 100% yield). $^1H$ NMR (400 MHz, $CDCl_3$) δ 3.35 (t, J = 6.7 Hz, 2H), 2.78 (t, J = 6.8 Hz, 2H), 1.70 (p, J = 6.8 Hz, 1H), 1.20 (bs, 1H).

**Synthesis of Compound 8.**[2] Compound **6** (1 g, 2.03 mmol) and **7** (0.406 g, 4.06 mmol) were stirred overnight at RT in DCM (10 mL). The reaction mixture was diluted with DCM (50 mL) and washed two times with 50 mL of saturated solution of $NaHCO_3$, two times with 50 mL 0.1 M HCl and 1 time with brine. The organic phase was dried ($MgSO_4$), filtrated and the filtrate concentrated *in vacuo* to obtain compound **8** as a pale yellow oil (0.83 g, 1.88 mmol, 90%). $^1H$ NMR (400 MHz, $CDCl_3$) δ 7.60 (s, 1H), 7.06 (s, 1H), 6.39 (q, J = 6.4 Hz, 1H), 4.31 – 4.09 (m, 6H), 3.97 (s, 3H), 3.42 (t, J = 6.6 Hz, 2H), 2.55 (t, J = 7.2 Hz, 2H), 2.20 (p, J = 6.8 Hz,2H), 1.97 – 1.90 (m, 2H), 1.69 (d, J = 6.4 Hz, 3H), 1.28 (t, J = 7.1 Hz, 3H).

**Synthesis of Compound 9.**[2] Compound **8** (0.83 g, 1.88 mmol) was dissolved in THF (10 mL). LiOH (0.1 g, 2.26 mmol) was dissolved in water (5 mL) and added dropwise to the mixture. The reaction mixture was left stirring at RT until TLC showed complete consumption of the starting material. Dowex® 50WX8 hydrogen form was added to the reaction mixture until neutral pH. The resin was filtered off and the filtrate was concentrated under reduced pressure to give compound **9** as a pale yellow oil (0.8 g, 1.88 mmol, 100%). $^1H$ NMR (600 MHz, $CDCl_3$) δ 7.58 (s, 1H), 6.98 (s, 1H), 6.37 (q, J = 6.4 Hz, 1H), 5.30 (s, 0H), 4.94 (t, J = 6.1 Hz, 1H), 4.12 (t, J = 6.2 Hz, 2H), 3.94 (s, 3H), 3.35 (t, J = 6.6 Hz, 2H), 3.24 (m, 2H), 2.60 (t, J = 7.1 Hz, 2H), 2.19 (p, J = 6.6 Hz, 2H), 1.77 (q, J = 6.6 Hz, 2H), 1.59 (d, J = 6.4 Hz, 3H).

**Synthesis of PC-CL (2).** Compound **9** (56 mg, 0.13 mmol) was transferred into a glass vial and dissolved in anhydrous DMF (0.5 mL). DIPEA (76.5 µL, 0.45 mmol, 9 eq), HOAt (16 mg, 0.12 mmol, 2.5 eq) and HATU (46 mg, 0.12 mmol, 2.5 eq) were added to the vial and then after everything was completely dissolved, $NH_2$-PEG(1kDa)-$NH_2$ (50 mg, 0.05 mmol) was added. The reaction was left stirring overnight at RT. The reaction mixture was loaded on a Sephadex LH-20 gel size exclusion column which was eluted with MeOH/DCM (1:1). The fractions containing product were combined and concentrated under reduced pressure to give photocleavable crosslinker **9** as a yellow oil (35 mg, 0.02 mmol). $^{1}H$ NMR (600 MHz, $CDCl_3$) δ 7.57 (s, 2H), 6.99 (s, 2H), 6.36 (q, J = 6.4 Hz, 2H), 5.02 (t, J = 6.1 Hz, 2H), 4.10 (t, J = 6.2 Hz, 4H), 3.95 (s, 6H), 3.69 – 3.59 (m, 82H), 3.56 (t, J = 5.1 Hz, 4H), 3.49 – 3.44 (m, 4H), 3.28 – 3.20 (m, 3H), 1.78 (q, J = 6.6 Hz, 3H), 1.59 (d, J = 6.5 Hz, 6H). $^{13}C$ NMR (151 MHz, $CDCl_3$) δ 172.89, 155.37, 153.92, 147.16, 139.76, 133.89, 109.17, 108.12, 70.50 (d, J = 3.1 Hz), 70.18, 69.73, 68.98, 68.51, 56.34, 49.18, 39.47, 38.61, 32.56, 29.04, 24.96, 22.24.

**Hyaluronic Acid Functionalization with BCN-$NH_2$ (1).** Hyaluronic acid 60 kDa (400 mg, 1 mmol of disaccharide units), purchased from Lifecore Biomedical (Chaska, MN, USA) was dissolved in 40 mL of MES buffer (100 mM, pH=6). 4-(4,6-dimethoxy-1,3,5-triazin-2-yl)-4-methyl-morpholinium chloride (DMTMM) (280 mg, 1 mmol) was added to the hyaluronic acid solution and left stirring until full dissolution. BCN-$NH_2$ (**5**) (50 mg, 0.150 mmol) was dissolved in 5 mL of ACN/$H_2O$ (2:1) and added dropwise to the hyaluronic acid solution. The reaction mixture was left stirring overnight at RT. The mixture was transferred in a 10 kDa cutoff dialysis bag and dialyzed first against 10% ACN/$H_2O$ solution for 24 h, then against 0.1 M $NaHCO_3$ solution for another 24 h and finally against mQ water for the final 24 h. The solution was then freeze dried to obtain a fluffy white solid of functionalized HA with a 15% degree of functionalization of BCN (**1**) (420 mg, 100%).

#### 4) SPPS of Peptides 3 ($N_3$-GGRGDSP) and 4 (AcK($N_3$)RDIPVSLRSGDRK($N_3$))

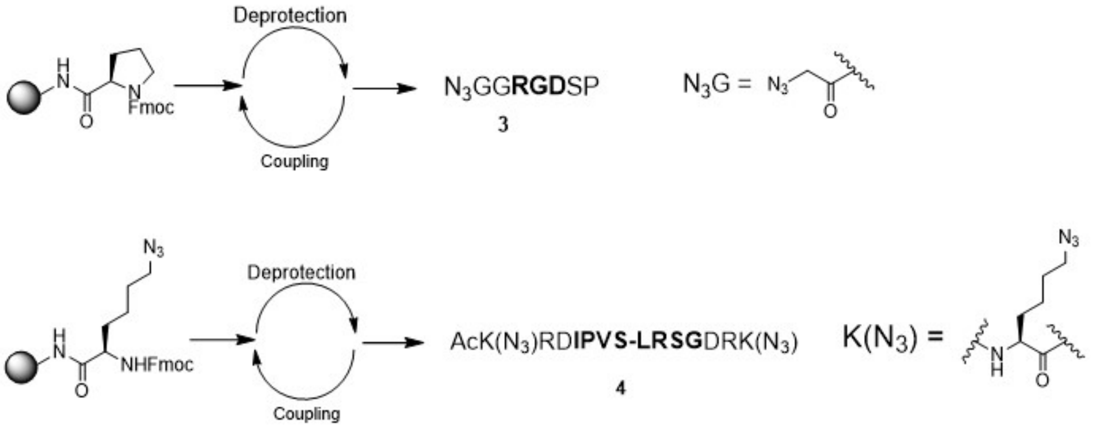


**Scheme S3.** General scheme for the SPPS of compounds **3** and **4**.

#### Synthesis of Peptide 3

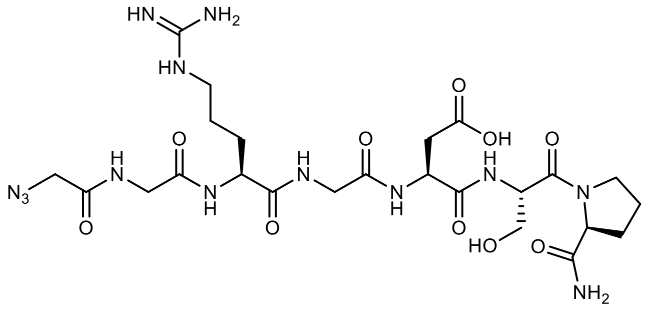

The resin-bound peptide $N_3$-GGRGDSP-$NH_2$ was synthesized by Fmoc SPPS on Rink amide resin (0.25 mmol scale). The resin was washed with DMF (3x) and dichloromethane (DCM, 3x) prior to peptide cleavage/deprotection (95:5 TFA:$H_2O$, 20 mL, 2 h) and precipitation (diethyl ether, 180 mL, 0 °C, 2x). The crude peptide was purified via RP-HPLC using a 55-minute gradient from 5-100% acetonitrile:$H_2O$; lyophilization yielded the final product ($N_3$-GGRGDSP-$NH_2$) as a white solid (120 mg, 0.18 mmol, 72% overall yield). MALDI-TOF MS: calculated for $C_{24}H_{39}N_{13}O_{10}^+$ $[M + 1H]^+$, e/z 670.29; observed 670.38.

**Synthesis of Peptide 4**

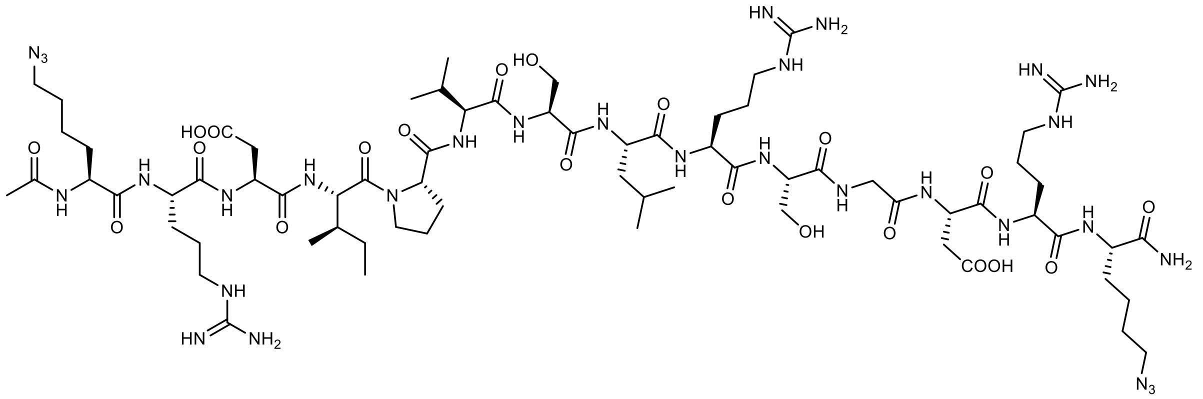


The resin-bound peptide Ac-K($N_3$)RDIPVS-LRSGDRK($N_3$)-$NH_2$ was synthesized by Fmoc SPPS on a Rink amide resin (0.25 mmol scale) and capped with an acetyl moiety using a solution of acetic anhydride 10% DIPEA 5% and DMF 85% (v/v/v). The resin was washed with DMF (3x) and dichloromethane (DCM, 3x) prior peptide cleavage/deprotection (95:5 TFA:$H_2O$, 20 mL, 2 h) and precipitation (diethyl ether, 180 mL, 0 °C, 2x). The crude peptide was purified via RP-HPLC using a 55-min gradient from 5-100% acetonitrile:$H_2O$; lyophilization yielded the final product (Ac-K($N_3$)RDIPVS-LRSGDRK($N_3$)-$NH_2$) as a white solid (120 mg, 0.18 mmol, 72% overall yield). MALDI-TOF: calculated for $C_{24}H_{39}N_{13}O_{10}^+$ [M + 1H]+, 1719.95; observed e/z 1719.48

**5) References**

(1) Dommerholt, J.; Schmidt, S.; Temming, R.; Hendriks, L. J.; Rutjes, F. P.; van Hest, J. C.; Lefeber, D. J.; Friedl, P.; van Delft, F. L. Readily accessible bicyclononynes for bioorthogonal labeling and three-dimensional imaging of living cells. *Angew. Chem. Int. Ed.* **2010**, *49* (49), 9422-9425.

(2) LeValley, P. J.; Neelarapu, R.; Sutherland, B. P.; Dasgupta, S.; Kloxin, C. J.; Kloxin, A. M. Photolabile linkers: Exploiting labile bond chemistry to control mode and rate of hydrogel degradation and protein release. *J. Am. Chem. Soc.* **2020**, *142* (10), 4671-4679.

### 6) NMR Spectra

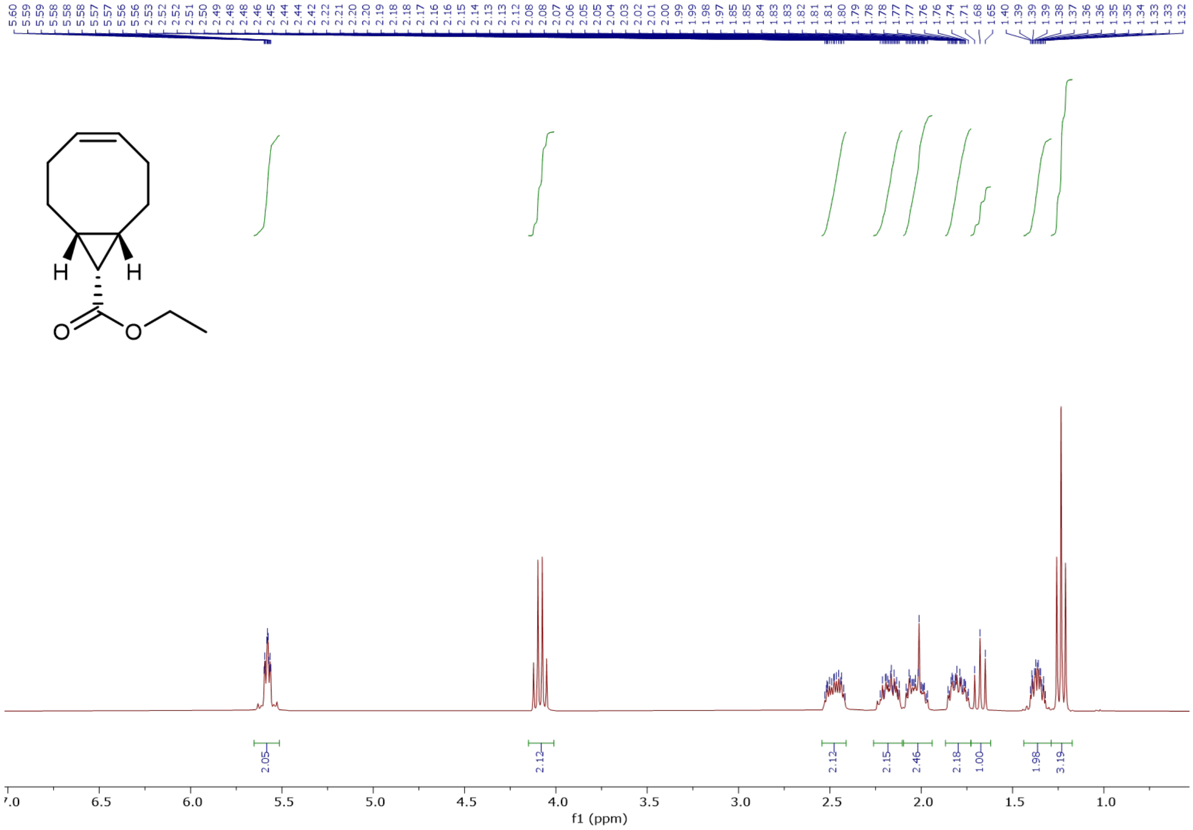


**NMR S1.** $^{1}$H NMR of compound **10**.

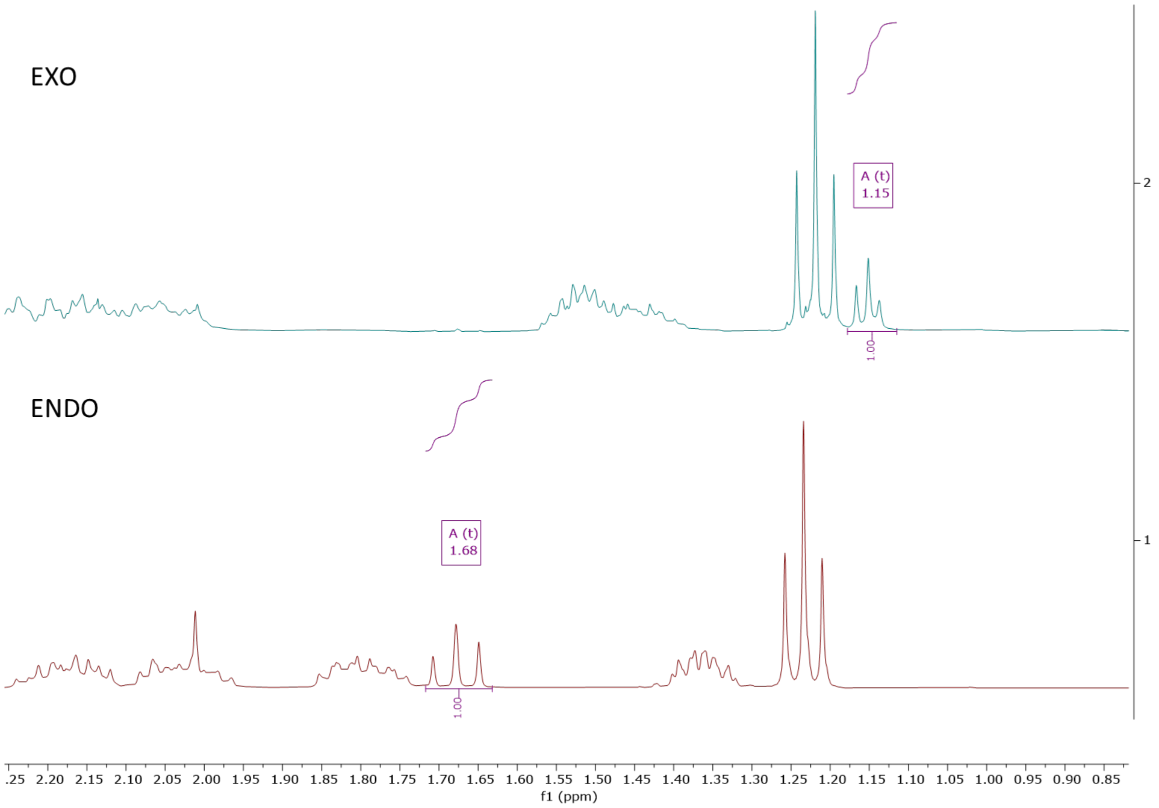


**NMR S2.** $^{1}$H NMR Comparison of Endo and Exo diastereomers of compound **10**.

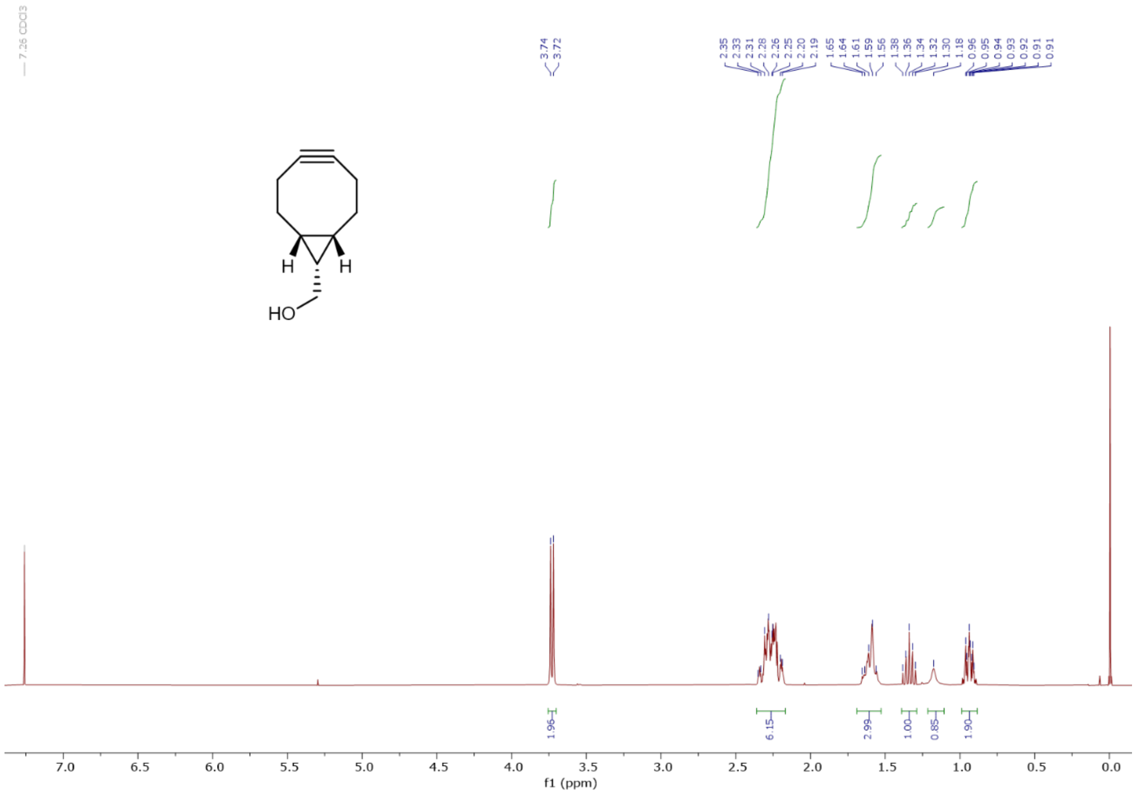

**NMR S3.** $^{1}$H NMR of compound **11**.

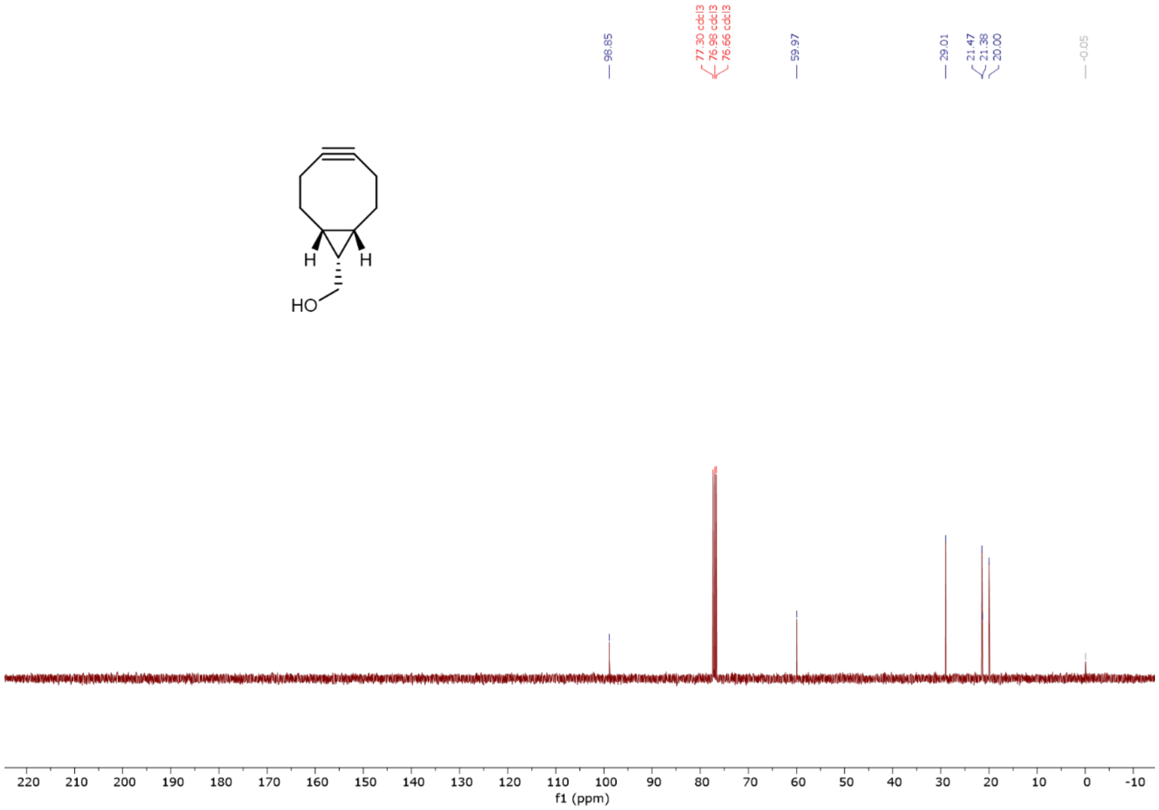

**NMR S4.** $^{13}$C NMR of compound **11**.

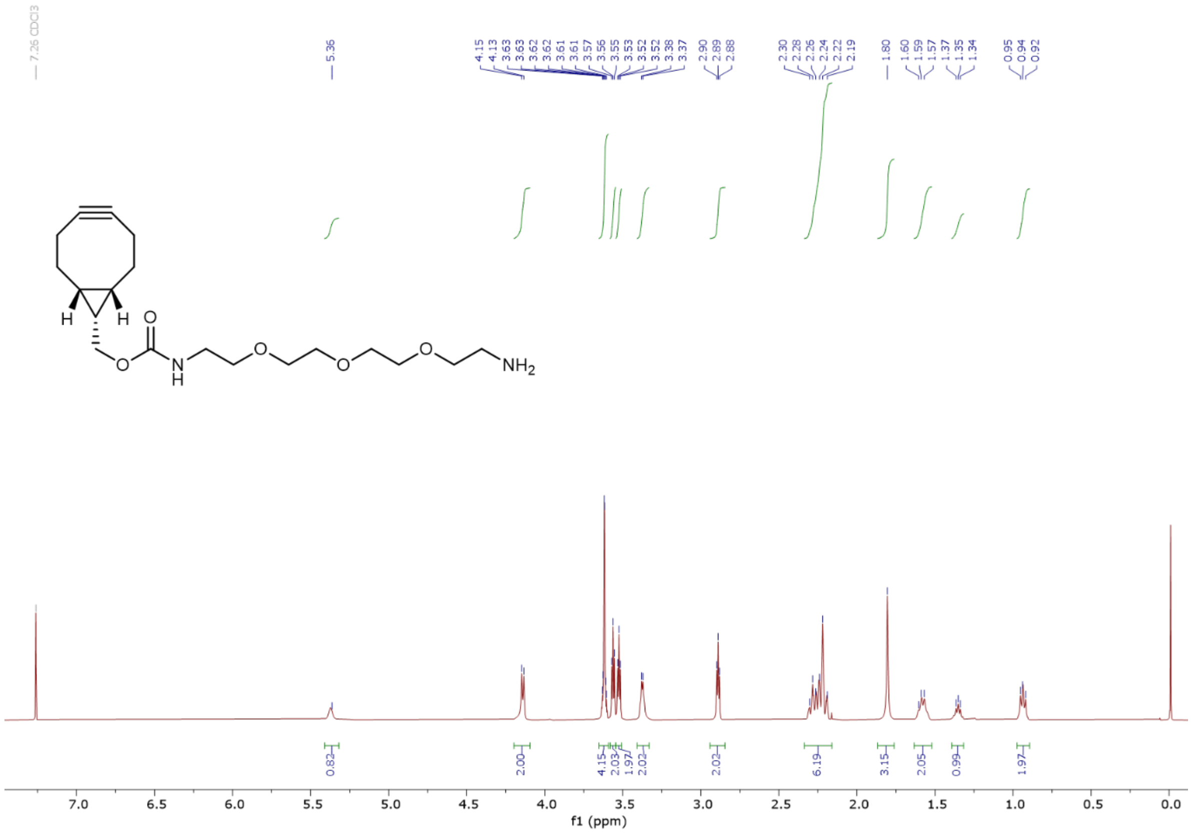


**NMR S5.** $^{1}$H NMR of compound **5**.

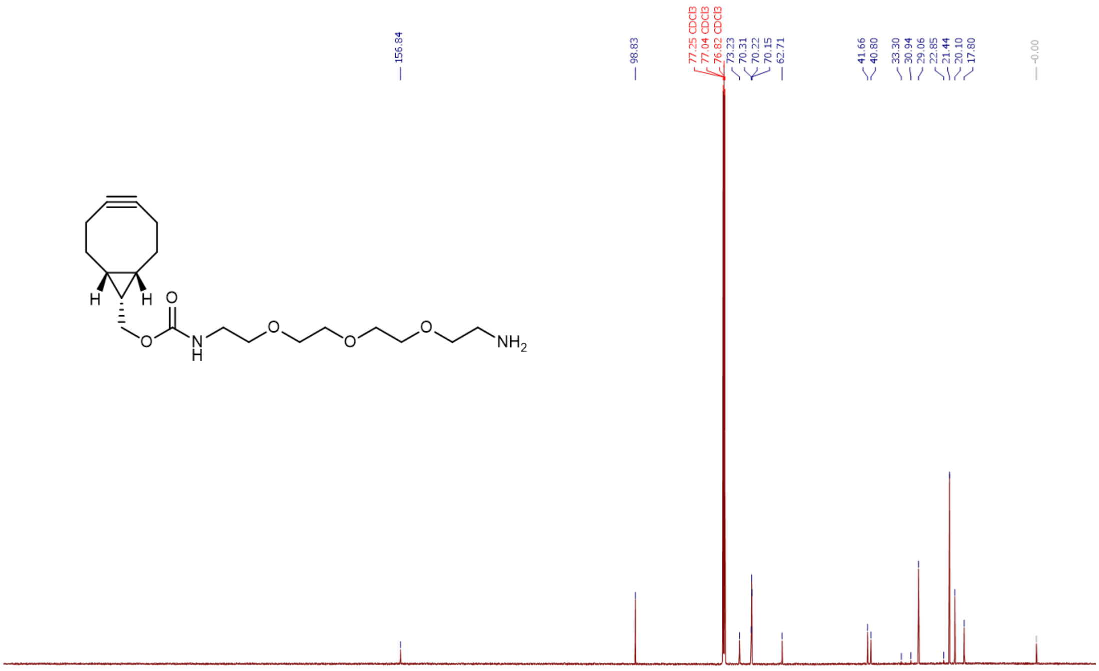


**NMR S6.** $^{13}$C NMR of compound **5**.

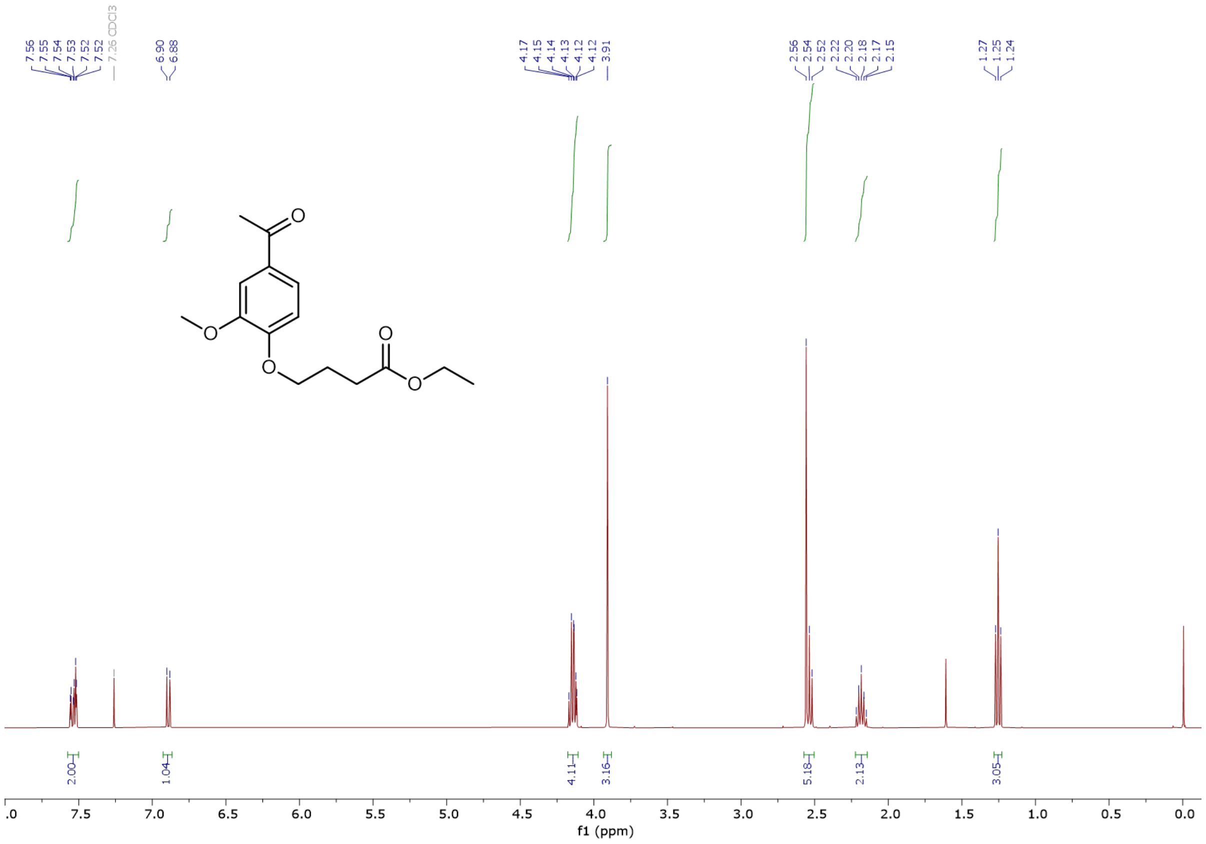

**NMR S7.** $^{1}$H NMR of compound **12**.

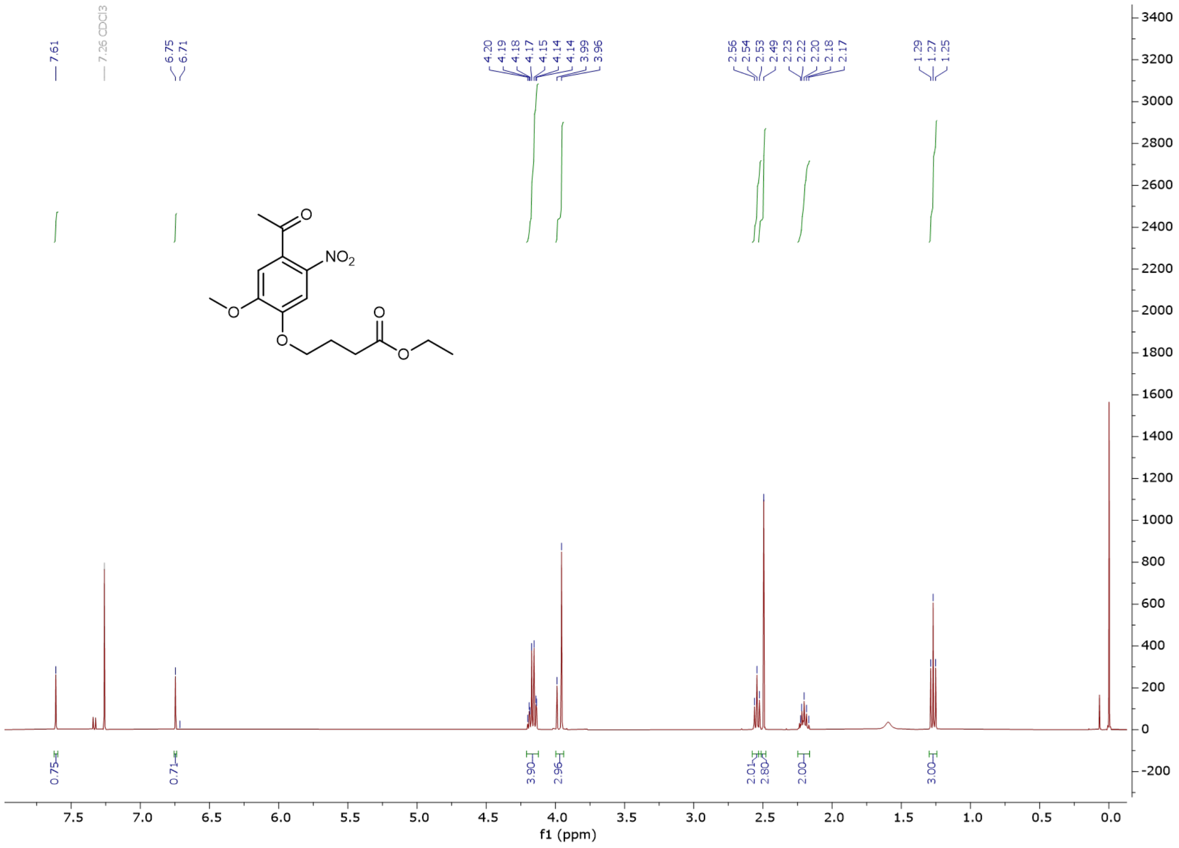

**NMR S8.** $^{1}$H NMR of compound **13**.

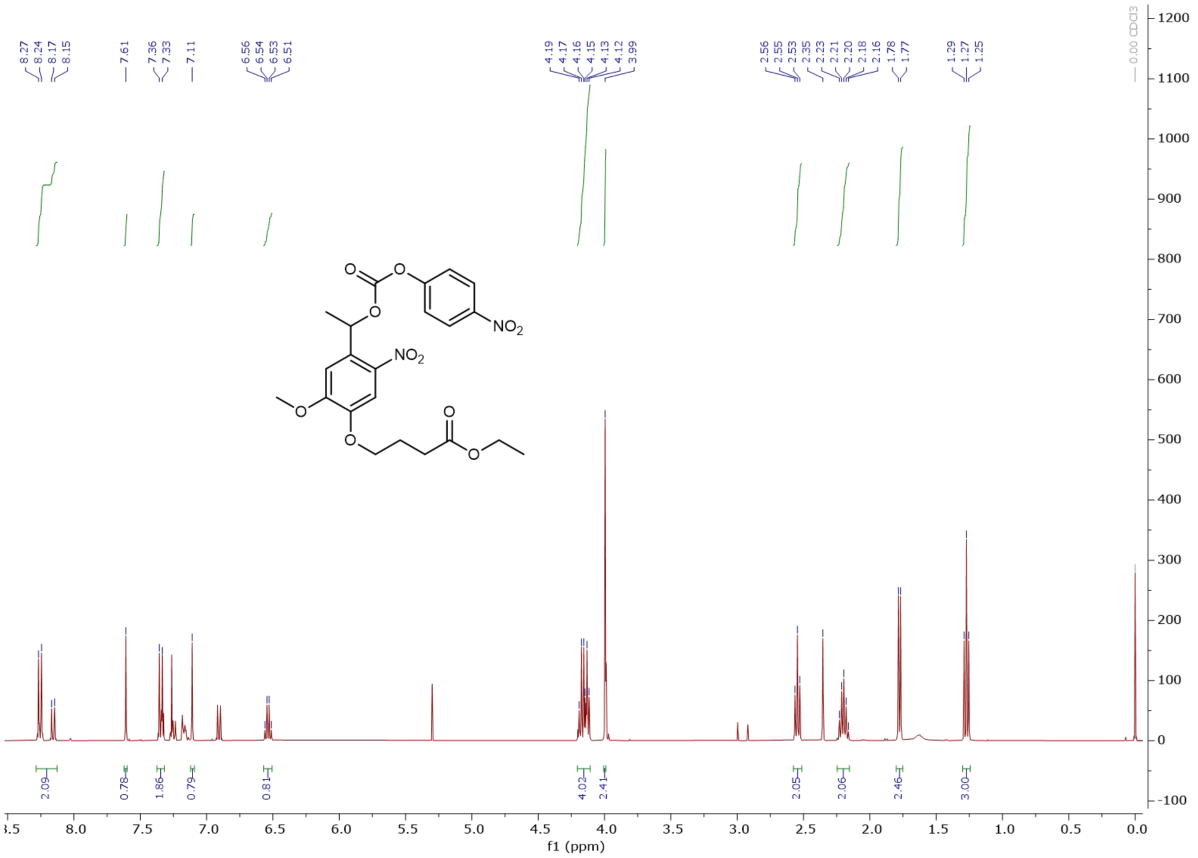

**NMR S9.** $^{1}$H NMR of compound **6**.

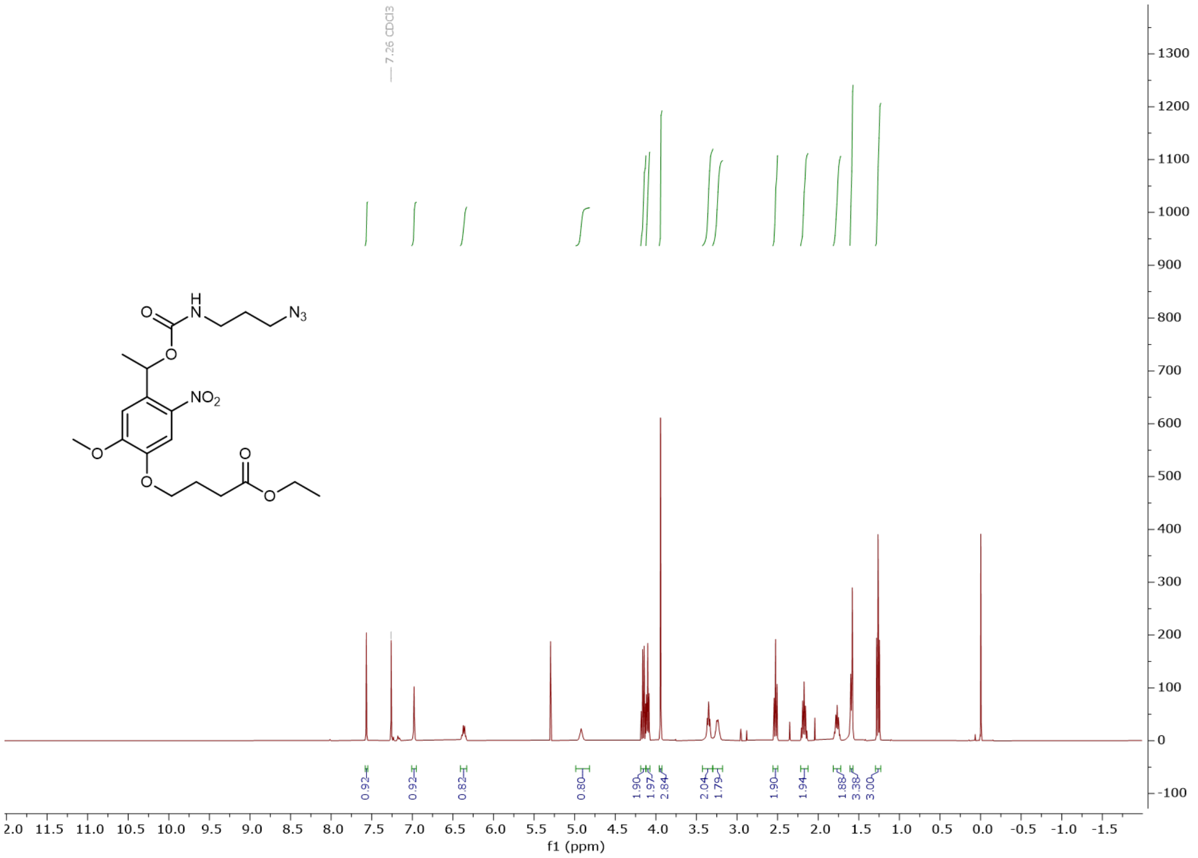

**NMR S10.** $^{1}$H NMR of compound **8**.

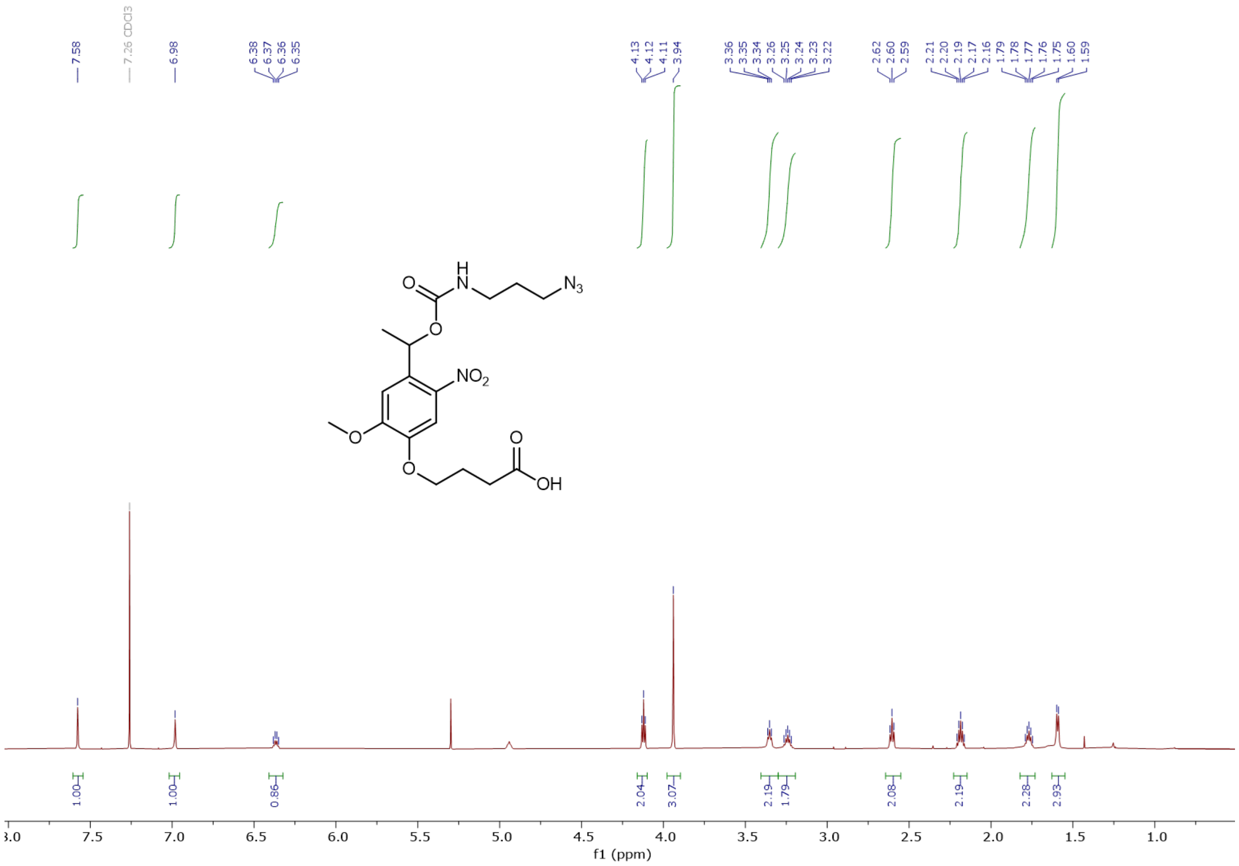

**NMR S11.** $^{1}$H NMR of compound **9**.

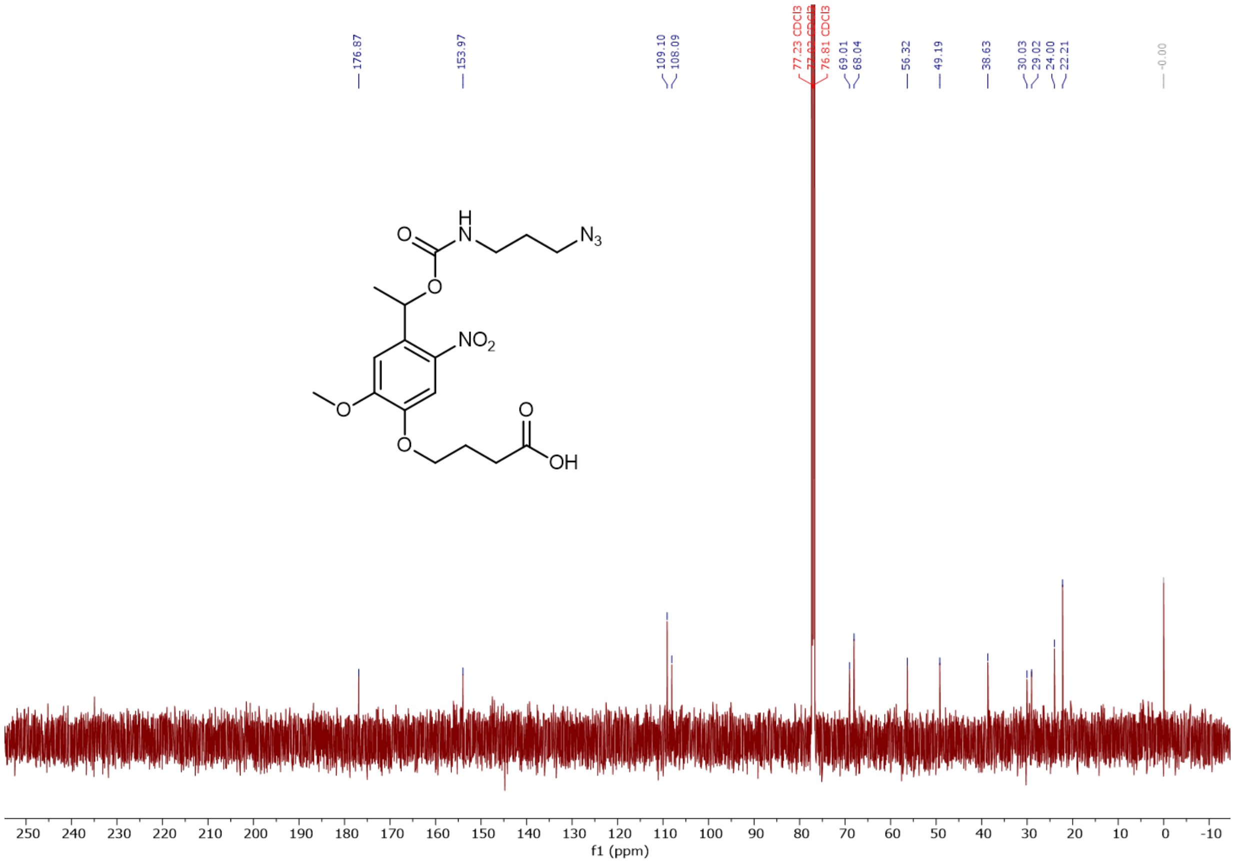

**NMR S12.** $^{13}$C NMR of compound **9**.

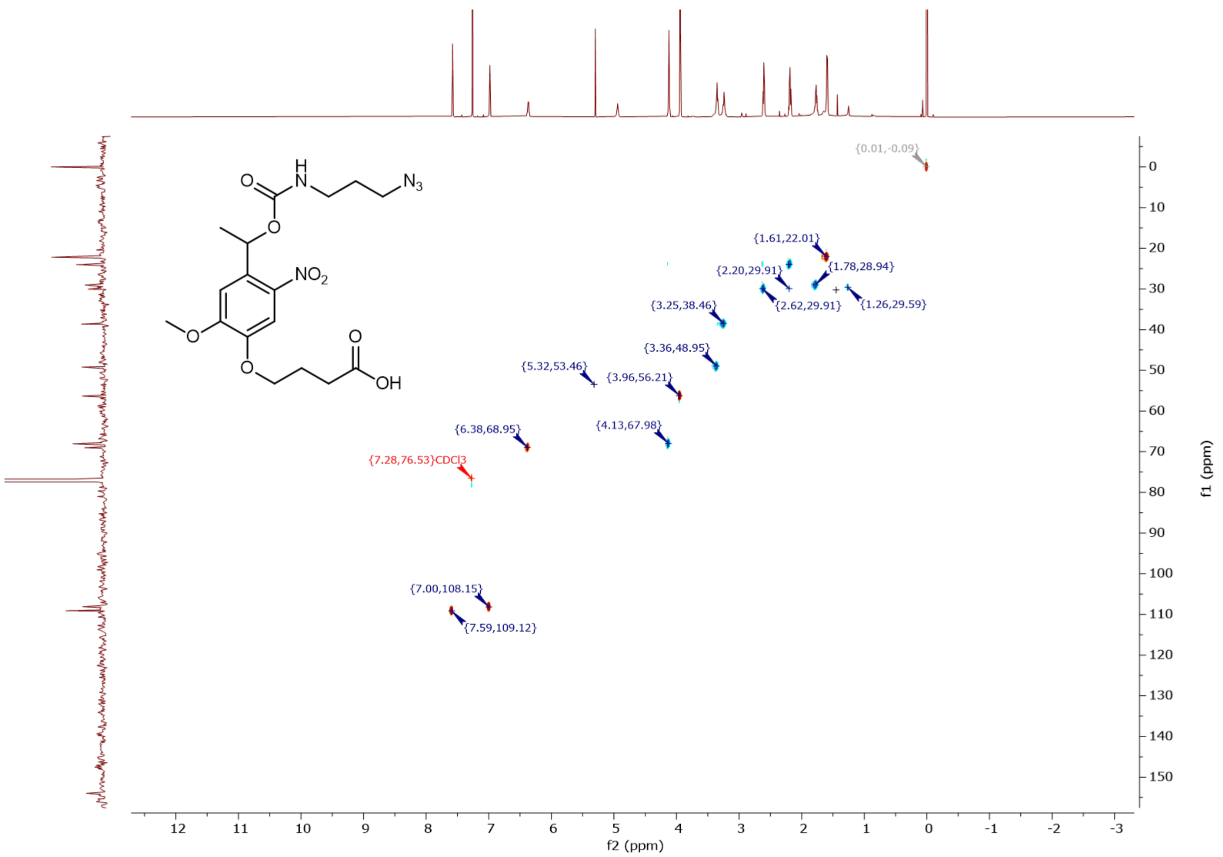

**NMR S13.** HSQC NMR of compound **9**.

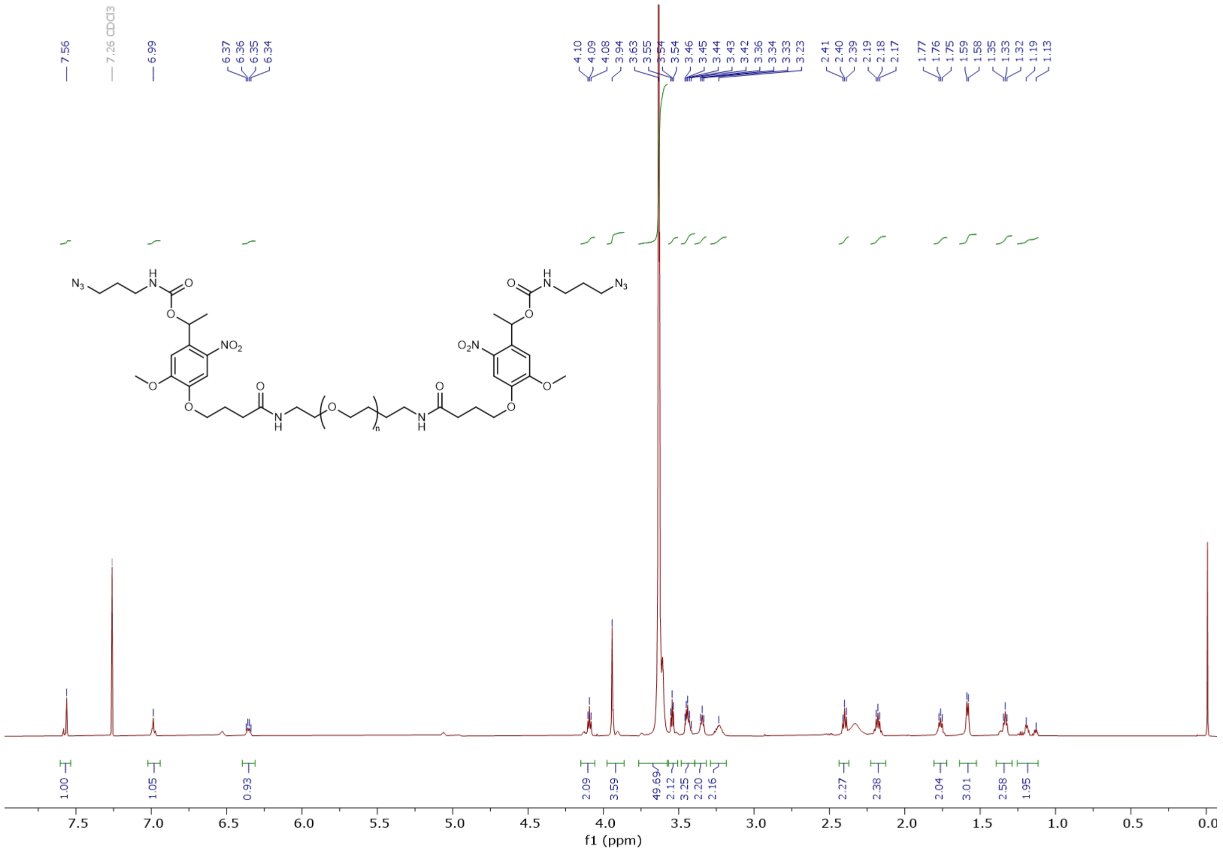

**NMR S14.** $^{1}$H NMR of compound **2**.

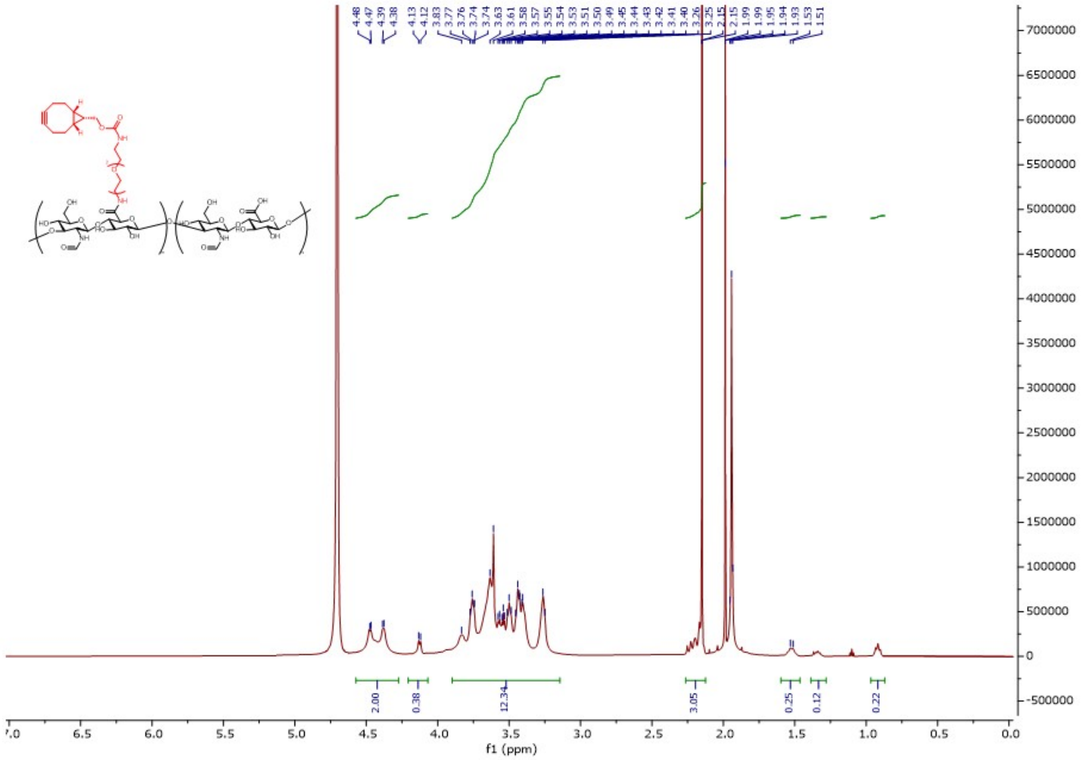


**NMR S15.** $^{1}$H NMR of compound **1** Anomeric protons at δ 4,47 – 4,37 (2H), BCN proton at δ 4,12 (2H 15%) endocyclic protons (3,90 -3,14) (12H) and NAc δ 2,15 (3H).

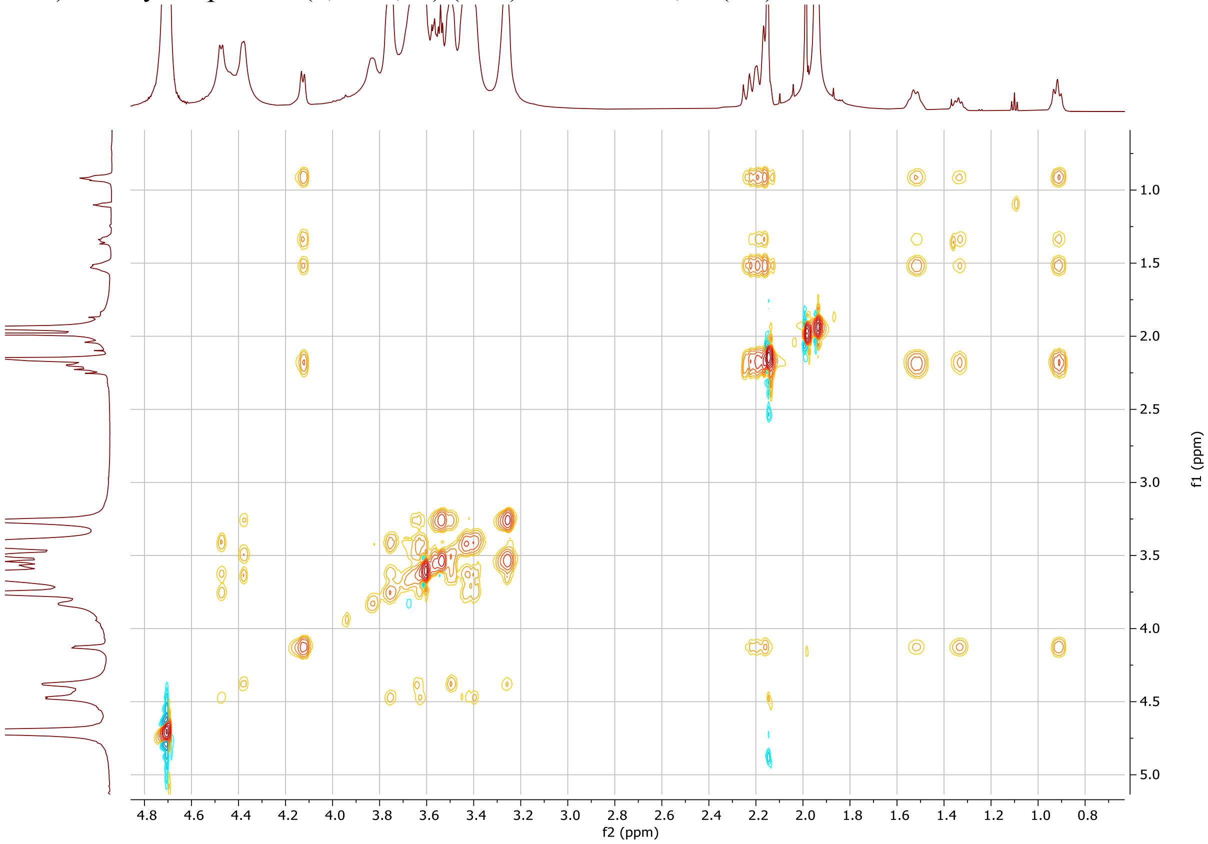


**NMR S16.** TOCSY NMR of compound **1**.

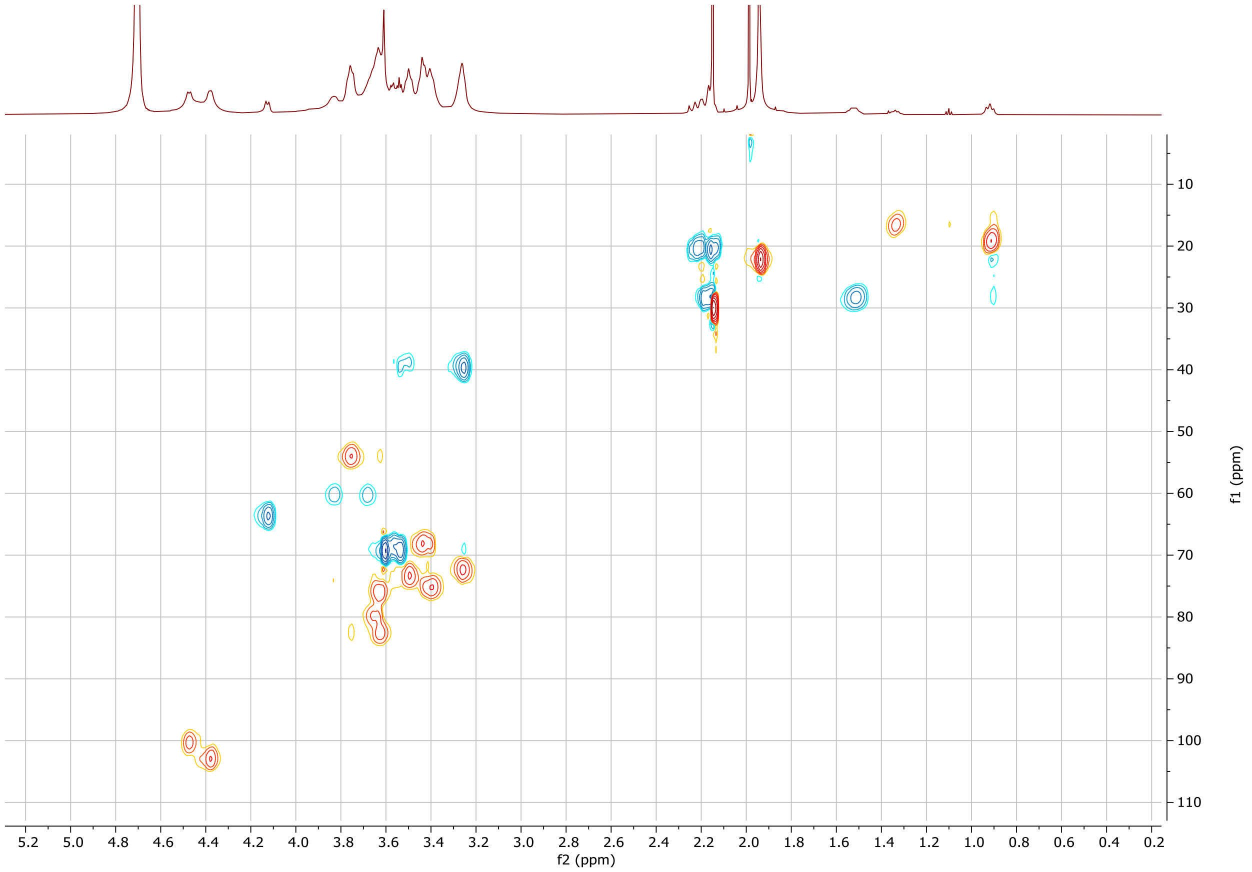


**NMR S17.** HSQC NMR of compound **1**.
